# Reshaping Organellar Translation and tRNA Metabolism: The Consequences of Photosynthesis Loss and Massive Horizontal Gene Transfer

**DOI:** 10.64898/2026.01.09.698701

**Authors:** Luis Federico Ceriotti, Leonardo M. Gatica-Soria, Kasavajhala V.S.K. Prasad, Rachael A. DeTar, Jessica M. Warren, Estefania Eichler, Joanna M. Chustecki, Christian Elowsky, Alan C. Christensen, Renchao Zhou, Daniel B. Sloan, M. Virginia Sanchez-Puerta

## Abstract

The transition to holoparasitism in plants precipitates the loss of photosynthesis, fundamentally altering the selective landscape acting on organellar genomes. These changes raise questions about the mechanisms by which the essential, coevolved machinery of translation responds to extreme genomic erosion and metabolic dependency. Integrating comparative genomics, tRNA sequencing, and subcellular localization assays, we elucidate the extensive rewiring of organellar translation systems and the tRNA-dependent tetrapyrrole biosynthesis pathway in the holoparasitic angiosperm family Balanophoraceae, which exhibits extreme reduction of tRNA content in plastid and mitochondrial genomes. We identified a rare evolutionary event: the putative intracellular transfer of the plastid initiator tRNA (tRNA-iMet) to the nucleus, which compensates for its loss from the plastid genome. We also demonstrate that the unusual UAG-to-Trp reassignment in the *Balanophora* plastid genetic code is driven by the loss of release factor pRF1 and the recruitment of a mutated nuclear tRNA-Trp. Furthermore, we reveal that the retention of organellar nuclear-encoded aminoacyl-tRNA synthetases is dictated by the presence/absence of cognate organellar tRNAs, which appear to be functional regardless of their foreign (horizontal transfer from the host plant) or native origins. Finally, we uncover a striking evolutionary asymmetry in nuclear-encoded ribosomal proteins: while plastid subunits exhibit elevated substitution rates consistent with relaxed selection and compensatory coevolution, mitochondrial subunits display high sequence conservation, likely maintaining compatibility with the extensive horizontal gene transfer observed in this lineage. Collectively, these findings represent some of the most extreme changes ever identified in the anciently conserved machinery of plant organellar translation.

## INTRODUCTION

The transition to a holoparasitic lifestyle represents one of the most radical evolutionary shifts in plants. This conversion precipitates the loss of photosynthesis, an event that fundamentally alters the selective landscape acting on organellar (plastid and mitochondrial) genomes. Recent findings suggest that this metabolic shift drives an extensive rewiring of plant organellar translation systems, characterized by the loss and functional replacement of aminoacyl-tRNA synthetases (DeTar, et al. 2024) and dramatic structural changes in plastid genomes (plastomes), including gene reduction, extreme AT nucleotide bias, and elevated substitution rates (Bellot and Renner 2016; Wicke and Naumann 2018; Su, et al. 2019). We hypothesize that the rigorous translational and metabolic demands of photosynthesis anchors the redundancy of organellar and cytosolic translation machinery (Raven 1995; Li, et al. 2017; Heinemann, et al. 2020). Relaxing this purifying selection likely triggers an evolutionary cascade that reshapes the plastid translation system and exerts correlated effects on mitochondrial translation. The magnitude of these effects may depend on how much translational machinery the organelles share.

Among holoparasites, the family Balanophoraceae stands as a paradigm of extreme genomic reduction and reorganization. Beyond typical reductive trends, this lineage displays unique genomic traits, including unprecedented deviations in the plastid genetic code and the extensive horizontal acquisition of tRNA and ribosomal protein genes in the mitochondrial genome (Svetlikova, et al.; Sanchez-Puerta, et al. 2017; Su, et al. 2019; Ceriotti, et al. 2021). However, a critical knowledge gap remains regarding the mechanistic consequences of these alterations: How do such profound genomic alterations affect tRNA metabolism and the functional integrity of organellar translation systems?

In the plastid, comparative analysis of hemiparasitic Santalales suggests that the transition to holoparasitism precipitated massive tRNA gene losses, concomitant with other drastic genomic reductions (Su, et al. 2019; Kim, et al. 2023; Ceriotti, et al. 2025). Currently, the gene encoding tRNA-Glu (*trnE*) is the sole tRNA retained in the plastome of most Balanophoraceae species (Su, et al. 2019; Kim, et al. 2023). This specific retention is attributed to its dual role in tetrapyrrole biosynthesis and translation (Barbrook, et al. 2006). Tetrapyrrole synthesis produces chlorophyll and heme. Although chlorophyll is not required in the absence of photosynthesis, heme is necessary for multiple cellular functions, such as mitochondrial respiration, underscoring its fundamental importance regardless of trophic mode (Tanaka and Tanaka 2007). However, even this critical gene is absent in species of *Lophophytum* and *Ombrophytum*, which carry no plastid-encoded tRNAs (Ceriotti, et al. 2021; Ceriotti, et al. 2025). In the genus *Balanophora*, the only retained tRNA gene in the plastome is tRNA-Glu, but it is aberrant and likely to function exclusively in the tetrapyrrole biosynthesis pathway rather than in plastid translation (Su, et al. 2019). Comparisons across the family Balanophoraceae provide the opportunity to catch radical transitions in tetrapyrrole biosynthesis and plastid-nuclear interactions “in the act”.

This systematic erosion of the plastid translation machinery, including the loss of plastid-encoded tRNAs and the corresponding organellar aminoacyl-tRNA synthetases (aaRSs), necessitates functional replacement. To date, there has been only one documented case of a functional transfer of an organellar tRNA gene to the nuclear genome with subsequent targeting back to the organelle (Berrissou, et al. 2024), suggesting that this specific route of replacement is either evolutionarily rare or remains significantly underexplored. Instead, the import of cytosolic counterparts (nuclear-encoded tRNAs and the retargeting of cytosolic aaRSs to the organelles) appears to be the dominant mechanism employed by plants to compensate for the functional loss of native organellar genes (Salinas-Giegé, et al. 2015; Warren and Sloan 2020).

Parallel to these intracellular shifts, plant mitochondrial genomes are characterized by frequent acquisitions of foreign tRNA genes via horizontal gene transfer (HGT). While many insertions may remain non-functional, there are demonstrated cases where foreign tRNAs of plastid (Joyce and Gray 1989; Marchfelder, et al. 1990), bacterial (Kitazaki, et al. 2011; Knie, et al. 2015), and fungal origin (Sinn and Barrett 2020; Warren, et al. 2025) are expressed and active within the recipient mitochondria. This phenomenon is particularly pronounced in the Balanophoraceae, where host-to-parasite transfer reshapes the mitochondrial genome (Sanchez-Puerta, et al. 2019; Roulet, et al. 2020; Garcia, et al. 2021). This family exhibits a remarkable spectrum of tRNA replacement: *Lophophytum* spp. have replaced all native mitochondrial tRNAs with six to eight foreign homologs acquired from Fabaceae host plants (Sanchez-Puerta, et al. 2019; Roulet, et al. 2025), while the closely related *Ombrophytum subterraneum* presents a mosaic state, possessing a highly incomplete complement of both native and foreign tRNA genes (Roulet, et al. 2020). Conversely, the mitochondrial genomes of *Balanophora yakushimensis* and *B. laxiflora* retain only a single native tRNA gene, known as *trnfM* or the tRNA-iMet gene (Yu, et al. 2025). However, despite this clear evidence of transfer and retention, the functionality of these specific host-derived genes and the potential co-transfer of their corresponding aaRSs remain unexplored.

Finally, the challenges of extreme heterotrophy extend to the assembly of organellar ribosomes, which requires the coordinated expression of subunits encoded by both nuclear and organellar genes, a balance maintained by cytonuclear coevolution (Burton and Barreto 2012; Weng, et al. 2016; Havird, et al. 2017; Tressel, et al. 2025). In Balanophoraceae, this interplay is threatened by the accelerated evolution of plastid ribosomal genes (Schelkunov, et al. 2019; Ceriotti, et al. 2025) and the replacement of native mitochondrial ribosomal proteins with foreign homologs (Sanchez-Puerta, et al. 2019; Roulet, et al. 2024), creating chimeric protein complexes of unknown stability (Sloan, et al. 2018).

In this study, we integrate comparative genomics, tRNA sequencing, and subcellular localization assays to investigate the translational machinery in Balanophoraceae. Specifically, we aim to: i) assess the functionality of native and foreign organellar tRNA genes via expression and CCA-tailing analysis; ii) characterize the genetic alterations underpinning plastid genetic code changes; iii) analyze the coevolutionary dynamics between organellar tRNA content and the retention of specific aaRSs; iv) determine the subcellular localization of tetrapyrrole biosynthesis enzymes in the absence of plastid-encoded tRNA-Glu; and v) investigate the subunit composition and cytonuclear evolution of organellar ribosomes.

## RESULTS

### Intracellular gene transfer of the plastid tRNA-iMet gene to the nucleus

The evolution of organellar genomes in Balanophoraceae is marked by a complex history of tRNA gene loss, HGT, and intracellular functional transfer to the nucleus (**Figure 1**). To investigate the expression and origin of the tRNA pool, we performed multiplex small RNA-seq (MSR-seq) on inflorescences of *Balanophora laxiflora*, *Lophophytum pyramidale*, and *Ombrophytum subterraneum*, using *Arabidopsis thaliana* as a control. Mapping statistics revealed a striking dominance of nuclear-encoded tRNAs in the holoparasites, accounting for >99.9% of mapped reads (**Table S1, Table S2**). This contrasts sharply with *Arabidopsis*, where only 65.6% of reads mapped to nuclear tRNAs, with significant contributions from plastid (32.3%) and mitochondrial (1.7%) genes (**Table S2**), consistent with previous reports (Ceriotti, et al. 2024). Analysis of 3′ termini revealed distinct patterns. While *B. laxiflora* displayed a high proportion of mature CCA-tailed tRNAs comparable to *Arabidopsis*, *L. pyramidale* and *O. subterraneum* exhibited elevated levels of CC-tailed transcripts (**Figure S1**). This likely reflects a higher degree of 3′-end degradation in these samples, potentially attributable to post-collection stability or the influence of species-specific secondary metabolites.

**Figure 1.**
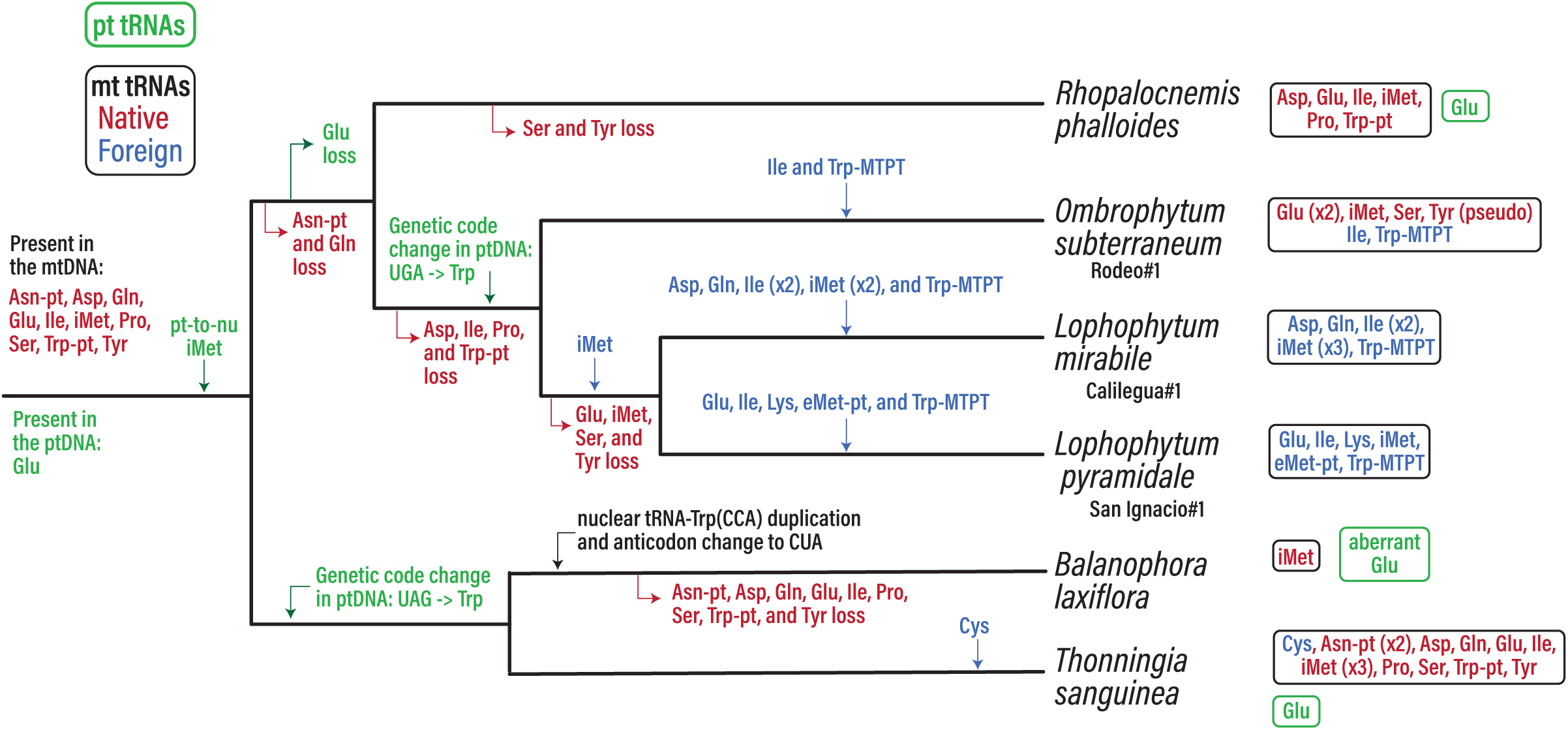
Evolution of organellar tRNAs in Balanophoraceae. The mitochondrial (mt), both foreign (in blue font) and native (in red font), and plastid (pt) tRNAs present in each organellar genome are shown on the far right. Only those Balanophoraceae for which plastid and mitochondrial genomes are available were included. The mitochondrial genomes have ancestrally acquired tRNA of plastid origin (pt) and plastid tRNAs embedded in plastid-derived mitochondrial DNA (MTPT).

Despite the overrepresentation of nuclear sequences, we also detected specific reads in *L. pyramidale* and *O. subterraneum* mapping to *Arabidopsis* plastid tRNA genes (**Table S3**, **Figure S2**). While most low-abundance hits likely represent background noise, the plastid initiator tRNA-iMet gene was consistently detected in *O. subterraneum* libraries (>80 reads), exceeding no-template controls (**Figure S2**). BLASTn analysis confirmed the plastid origin of these reads, which showed 92% identity to plastid tRNA-iMet gene from hemiparasitic Santalales and 87% to *Arabidopsis*, yet shared no homology with the assembled *O. subterraneum* mtDNA (available via https://github.com/lfceriotti/Balanophoraceae-MSRseq). This strongly implies a plastid-to-nucleus transfer of the tRNA-iMet gene.

To corroborate this intracellular gene transfer (IGT), we mined the nuclear genomes of *Balanophora fungosa* and *B. subcupularis* (Chen, et al. 2023) and identified a nuclear-encoded tRNA-iMet gene of clear plastid ancestry. These nuclear copies share 79% identity with the *Arabidopsis* plastid homolog and phylogenetically cluster with expressed transcripts from *B. laxiflora* (identified by mapping *B. laxiflora* MSR-seq data over *B. subcupularis* plastid tRNA-iMet; **Table 1, Table S4**) separate from the expressed tRNA reads of *O. subterraneum,* and of *L. pyramidale* (identified by mapping *L. pyramidale* MSR-seq data over *O. subterraneum* plastid tRNA-iMet; **Table S4)**. Although the tree topology **(Figure 2**) is not well resolved and could reflect either two independent transfers or an ancient intracellular transfer event from the plastid to the nucleus in the Balanophoraceae ancestor, the ancestral loss of the plastid-encoded tRNA-iMet gene from the Balanophoraceae plastomes suggests the latter scenario is more likely (**Figure 1**). Following this identification, we included the nuclear plastid-like tRNA-iMet sequences in our reference set for subsequent mapping analyses.

**Figure 2.**
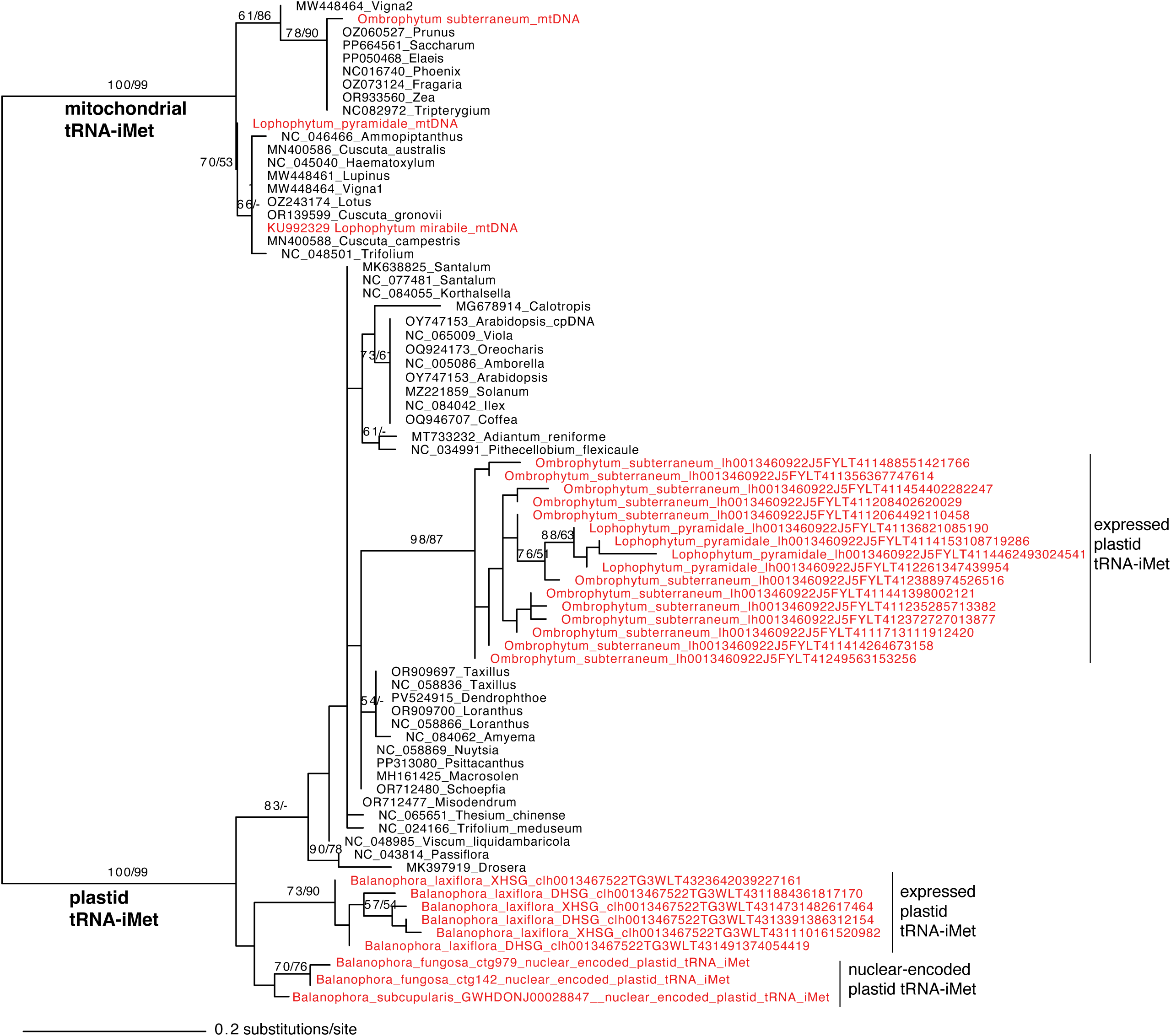
Maximum likelihood (ML) phylogenetic tree of the organellar tRNA-iMet across angiosperms. This tree includes mitochondrial and plastid-encoded genes and nuclear-encoded plastid tRNA-iMet genes identified in Balanophoraceae. Bootstrap support values >50% from ML (left) and maximum parsimony (right) analyses are shown above the branches.

**Table 1.** Number of CCA-tailed and total processed reads mapped to tRNAs of interest.

| Compartment | Reference tRNAs |  | *Balanophora laxiflora* XHSG |  | *Balanophora laxiflora* DHSG |  |
| --- | --- | --- | --- | --- | --- | --- |
|  | Species | Gene | CCA | Total | CCA | Total |
| mitochondria | *Balanophora laxiflora* | tRNA-iMet-CAT | 55 | 81 | 16 | 25 |
| nucleus | *Balanophora subcupularis* | plastid tRNA-iMet-CAT | 166 | 308 | 65 | 129 |
| nucleus | *Balanophora yakushimensis* | tRNA-Trp-CCA | 10672 | 17800 | 6266 | 10335 |
| nucleus | *Balanophora yakushimensis* | tRNA-Trp-CTA | 1044 | 1548 | 384 | 659 |
| plastid | *Balanophora laxiflora* | tRNA-Glu-TTC | 6 | 15 | 0 | 1* |
|  |  |  |  | ***Lophophytum pyramidale***<br>**SanIgnacio3** |  |  |
|  |  |  |  | **CCA** | **Total** |  |
| mitochondria | *Lophophytum* | tRNA-Glu-TTC (foreign) |  | 1450 | 3701 |  |
| mitochondria | *Lophophytum* | tRNA-Ile-CAT (foreign) ** |  | 120 | 457 |  |
| mitochondria | *Lophophytum* | tRNA-Lys-TTT (foreign) ** |  | 281 | 827 |  |
| mitochondria | *Lophophytum* | plastid tRNA-Trp-CCA (foreign) *** |  | 1238 | 2681 |  |
| mitochondria | *Lophophytum* | plastid tRNA-eMet-CAT (foreign) |  | 23 | 27 |  |
| mitochondria | *Lophophytum* | tRNA-iMet-CAT (foreign) |  | 2029 | 5126 |  |
| nucleus | *Lophophytum* | plastid tRNA-iMet-CAT |  | 1172 | 3512 |  |
|  |  |  |  | ***Ombrophytum subterraneum***<br>**Rodeo1** |  |  |
|  |  |  |  | **CCA** | **Total** |  |
| mitochondria | *Ombrophytum* | tRNA-Glu-TTC (Chr1) |  | 542 | 1493 |  |
| mitochondria | *Ombrophytum* | tRNA-Glu-TTC (Chr30) |  | 0 | 45 |  |
| mitochondria | *Ombrophytum* | tRNA-Ile-CAT (foreign) ** |  | 215 | 643 |  |
| mitochondria | *Ombrophytum* | tRNA-Ser-GGA |  | 0 | 0 |  |
| mitochondria | *Ombrophytum* | tRNA-Trp-CCA (foreign) *** |  | 1314 | 3095 |  |
| mitochondria | *Ombrophytum* | tRNA-Tyr-TTC (pseudogene?) ** |  | 222 | 1192 |  |
| mitochondria | *Ombrophytum* | tRNA-iMet-CAT |  | 4831 | 9169 |  |
| nucleus | *Ombrophytum* | plastid tRNA-iMet-CAT |  | 1180 | 6127 |  |

\* This read has a CC-tail
\*\* These genes have a genomically-encoded CCA-tail
\*\*\* These genes are within MTPTs.

### Expression and post-transcriptional processing of native and foreign organellar tRNAs

Transcriptomic profiling confirmed that mitochondrial tRNAs of diverse phylogenetic origins are expressed in the Balanophoraceae. Specifically, we detected transcripts for five of the six foreign tRNAs found in *L. pyramidale* mtDNA (including the tRNA-Trp gene within a foreign plastid-derived mitochondrial DNA, MTPT), and both foreign tRNAs resident in *O. subterraneum*. Conversely, transcriptomic evidence was absent for specific mitochondrial-encoded loci. The genes tRNA-eMet in *L. pyramidale* and tRNA-Glu (Chr30) and tRNA-Ser (Chr49) in *O. subterraneum* yielded negligible mapping reads (**Table 1)**. Furthermore, the few reads mapping to these genes also mapped equally well to other reference tRNAs, indicating these loci are likely not expressed and represent pseudogenes.

Post-transcriptional processing analysis revealed that all expressed mitochondrial tRNAs possess mature 3′-CCA tails. With the exception of a few genes where the CCA is genomically encoded (**Table 1**), these tails are added post-transcriptionally, confirming that these genes are not only transcribed but also undergo transcript maturation. This processing includes the tRNA-iMet gene in *Balanophora* (the sole tRNA gene remaining in its mitochondrial genome) and the aberrant plastid-encoded tRNA-Glu gene (**Table 1**).

Analysis of base modification profiles, inferred from reverse transcriptase-induced “hard-stop” signatures and/or breakpoints of tRNA-derived fragments (tRFs), revealed conserved patterns across the parasitic species (**Figure S3**). In both *L. pyramidale* and *O. subterraneum*, the mitochondrial tRNA-Ile and tRNA-iMet genes displayed pronounced hard-stop signals indicative of specific base modifications at conserved positions. Similarly, the tRNA-iMet gene of *B. laxiflora* exhibited a clear modification footprint. Remaining tRNAs showed negligible signals of base modification. For the aberrant plastid-encoded tRNA-Glu gene of *Balanophora*, low read coverage precluded a reliable assessment of its modification profile.

### Genomic and transcriptomic basis of plastid genetic code reassignment

Previous analyses identified two distinct genetic code deviations in Balanophoraceae plastomes: the reassignment of UAG to Trp in the *Balanophora* clade, and UGA to Trp in the *Lophophytum*+*Ombrophytum* clade (Su, et al. 2019; Ceriotti, et al. 2021). We investigated the molecular mechanism(s) facilitating these reassignments, focusing on the evolution of peptide release factors (RFs) and tRNA anticodons.

We hypothesized that reassignment requires the loss of the RF responsible for recognizing the reassigned stop codon. Consistent with the UAG-to-Trp change, we confirmed the absence of the plastid-targeted release factor pRF1 (specific for UAA and UAG) and the retention of pRF2 in the nuclear genomes of *Balanophora subcupularis* and *B. fungosa* (Chen, et al. 2023). Conversely, in *L. pyramidale* and *O. subterraneum* (UGA-to-Trp), we detected pRF1 but found no transcripts for pRF2 (specific for UAA and UGA), as previously observed for *L. mirabile* (Ceriotti, et al. 2021). By comparison, mitochondrial RFs (mRF1 and mRF2) are retained across all examined Balanophoraceae. These findings demonstrate that the loss of specific plastid RFs correlates with codon reassignment.

To identify the tRNA responsible for decoding UAG as Trp in *Balanophora*, we analyzed MSR-seq data. In *B. laxiflora*, we detected expressed and mature (CCA-tailed) transcripts of a tRNA-Trp carrying a mutated CUA anticodon (complementary to UAG), alongside the canonical CCA anticodon (**Table 1**). While reads mapping to the tRNA-Trp-CUA gene were less abundant than the canonical species, they showed clear evidence of post-transcriptional modification (**Figure S4**). To rule out the possibility of RNA editing as the source of the CUA anticodon, we mined the nuclear genomes of *B. fungosa* and *B. subcupularis* (Chen, et al. 2023). We identified a specific nuclear locus encoding tRNA-Trp-CUA, distinct from the canonical tRNA-Trp-CCA genes in the nuclear genomes of *B. subcupularis* and *B. fungosa* (**Figure 3A**). Further, phylogenetic analysis indicated that this suppressor tRNA originated from a duplication of the canonical nuclear gene followed by a mutation in the anticodon prior to *Balanophora* species diversification (**Figure 3B**).

**Figure 3.**
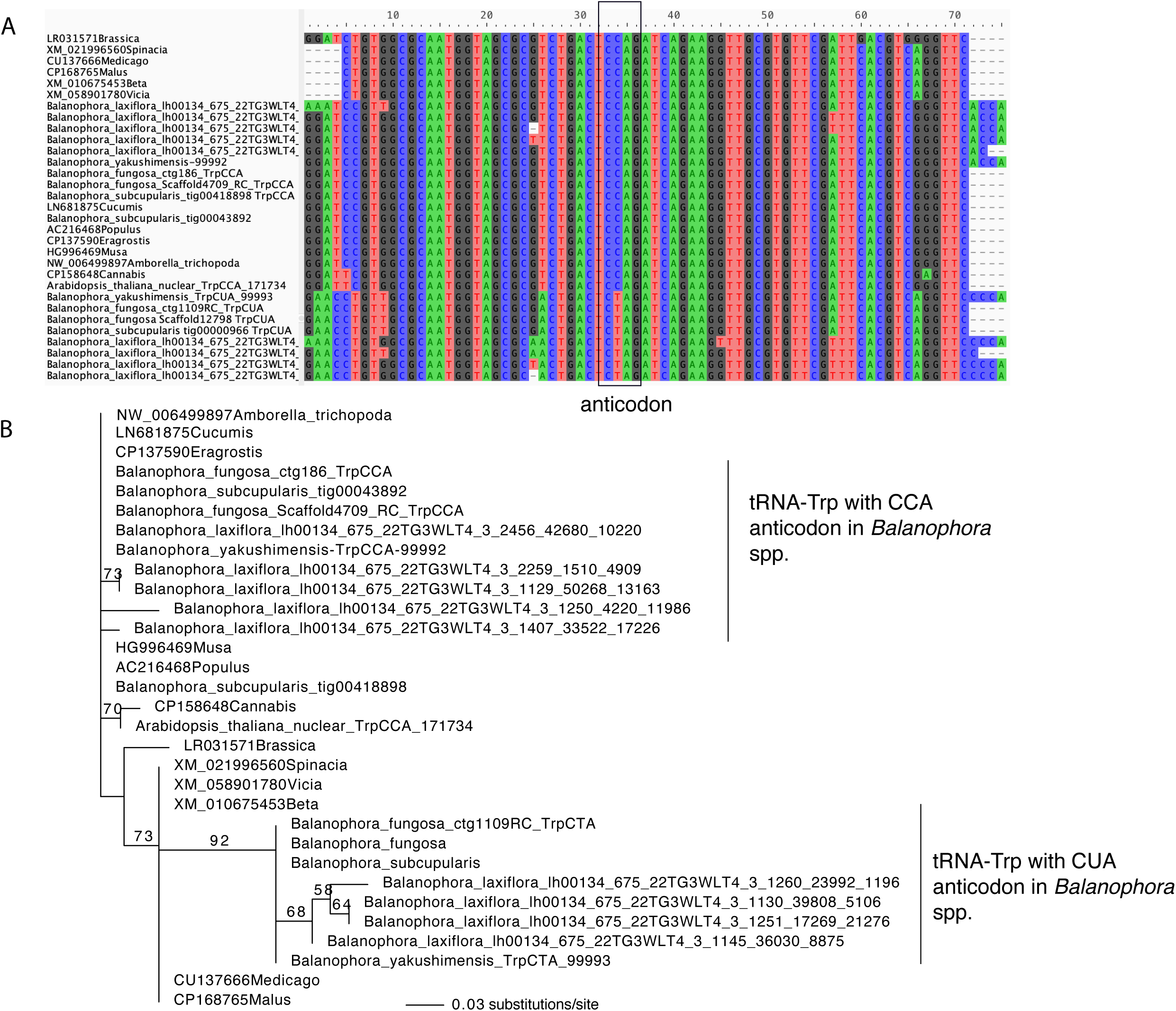
Nuclear-encoded tRNA-Trp in angiosperms. A. Nucleotide alignment of nuclear-encoded tRNA-Trp in angiosperms. B. Maximum likelihood phylogenetic tree of nuclear-encoded tRNA-Trp genes. Balanophora spp. present tRNA-Trp genes with anticodons CCA or CTA.

We tested whether the UGA-to-Trp reassignment in *Lophophytum* and *Ombrophytum* similarly involved a novel tRNA with a UCA anticodon. MSR-seq analysis revealed no evidence of a nuclear gene or post-transcriptional editing generating a UCA anticodon. While mapping against *Arabidopsis* reference genes revealed mismatches concentrated at position 36 of the anticodon loop, the anticodon triplet itself (positions 33–35) showed no modifications indicative of a C-to-U edit (**Figure S5**). Although we did detect rare reads with a UCA sequence, their relative abundance was negligible and comparable to background noise levels observed in *Arabidopsis* (**Table S5**), a species with a standard genetic code. Consequently, our data do not support the existence of a dedicated tRNA–Trp-UCA gene or editing event for UGA decoding in *Lophophytum* and *Ombrophytum.* Thus, it remains unclear how UGA is decoded to Trp in these species.

### Coevolution of organellar tRNA content and aaRS retention

The relaxation of selective pressure on photosynthesis typically drives the concurrent loss of plastid-encoded tRNAs and their cognate organellar aaRSs (Cai, et al. 2021; DeTar, et al. 2024). Given that *Balanophora* plastomes retain only the tRNA-Glu gene, while *Lophophytum* and *Ombrophytum* lack plastid tRNA genes entirely, we anticipated the loss of the complete organellar aaRS repertoire, with two functional exceptions: the organellar GluRS (required to charge the retained plastid tRNA-Glu in *Balanophora*) and the organellar PheRS (typically retained due to the two-subunit nature of the cytosolic PheRS) (Pett and Lavrov 2015; Warren, et al. 2023; DeTar, et al. 2024).

Our aaRS inventory corroborated the retention of these expected enzymes but also revealed a more complex landscape of retention and loss. In the majority of cases, the presence of an organellar aaRS mirrors the presence of expressed mitochondrial tRNAs, regardless of whether those tRNAs are native or foreign (**Table S6, Figure 4**). However, we identified three uncoupling events. In *L. pyramidale*, the foreign tRNA-Lys is actively expressed (with a genomically-encoded CCA-tail), yet the organellar LysRS is missing. The organellar HisRS and AsnRS are retained in *Lophophytyum/Ombrophytum* and *Balanophora*, respectively, despite the complete absence of organellar-encoded tRNAs (**Table S6**).

**Figure 4.**
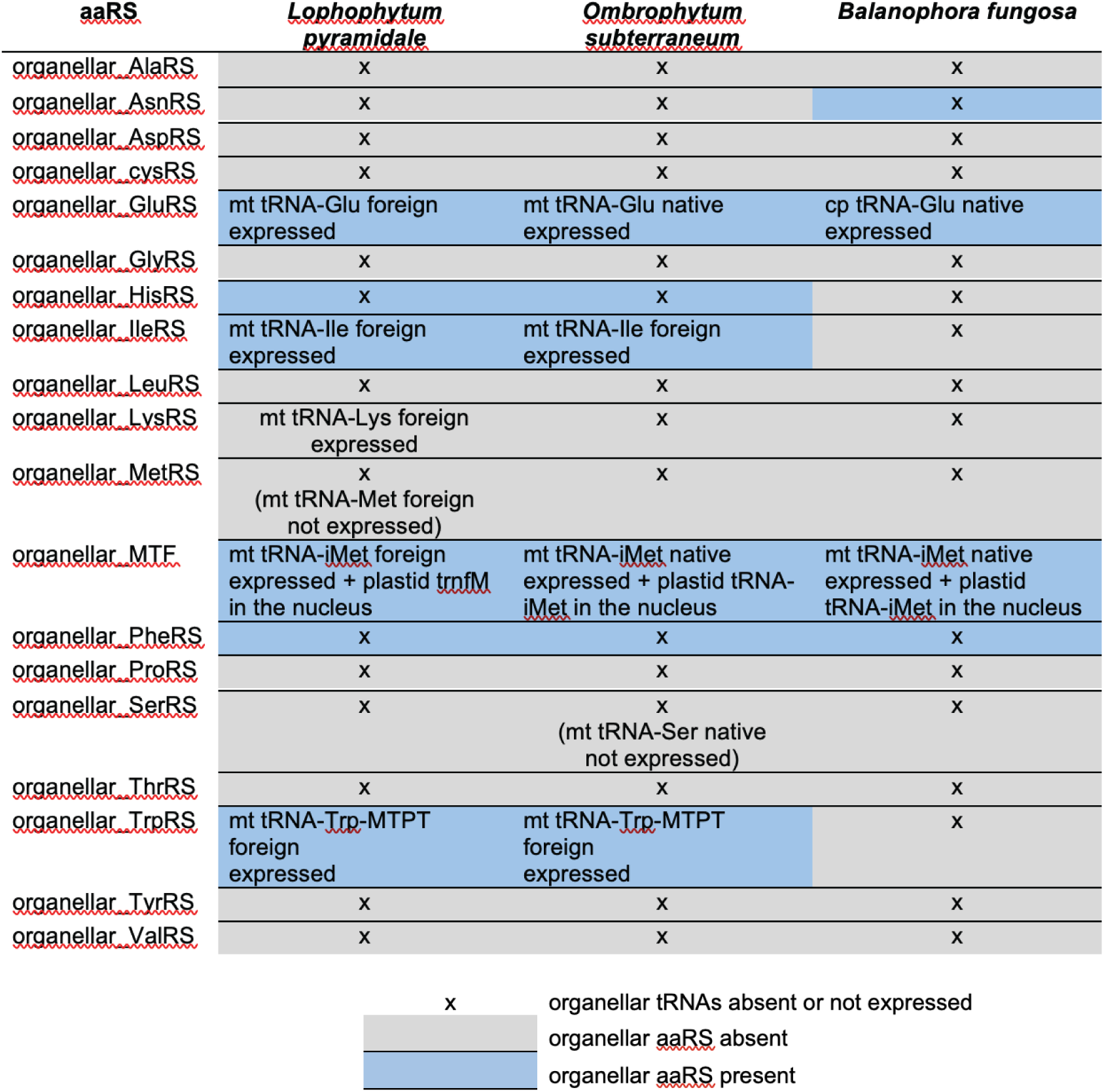
Co-retention of organellar aminoacyl-tRNA synthetases (aaRSs) and their cognate tRNAs in holoparasitic Balanophoraceae. The matrix illustrates the presence, absence, and origin of organellar aaRSs and their corresponding tRNAs across Lophophytum pyramidale, Ombrophytum subterraneum, and Balanophora spp.

We investigated whether the horizontal acquisition of foreign tRNAs in *Lophophytum* spp. and in *O. subterraneum* was accompanied by the co-transfer of their cognate aaRS genes. Phylogenetic reconstruction demonstrates that this is not the case: the aaRSs responsible for charging these foreign tRNAs are of native origin (**Figure S6**). Nearly the entire complement of cytosolic and organellar aaRSs remains native, with only two documented instances of HGT in the cytosolic compartment: the replacement of the native cytosolic ThrRS in *Balanophora* spp. by a foreign homolog and the acquisition of a foreign cytosolic TyrRS in *L. mirabile*, which coexists with the native copy (**Figure S6**).

The extensive loss of organellar aaRSs in Balanophoraceae necessitates the import of cytosolic enzymes to charge the remaining or imported tRNAs, a phenomenon observed in other non-photosynthetic lineages (DeTar, et al. 2024). This retargeting can evolve via gene duplication followed by neo-functionalization (acquisition of a transit peptide) or via alternative transcription/translation initiation of a single gene. We interrogated the high-quality nuclear genomes of *B. fungosa* and *B. subcupularis* (Chen, et al. 2023) for duplications of cytosolic aaRS genes. In all cases, we recovered only single-copy genes (**Table S6**). This absence of paralogs suggests that compensation is likely achieved through ambiguous targeting signals or alternative start sites within the single cytosolic locus, rather than through gene duplication.

### Subcellular localization of tetrapyrrole biosynthesis and the evolution of the plastid-encoded tRNA-Glu gene

The plastid-encoded tRNA-Glu gene, which is essential for tetrapyrrole biosynthesis, has undergone significant divergence (*Balanophora*) or complete loss (*Lophophytum + Ombrophytum*) in Balanophoraceae, while several lines of evidence suggest that tetrapyrrole biosynthesis persists in the plastids of this family (Ceriotti, et al. 2021; Chen, et al. 2023). However, it is unclear whether a plastid tRNA-Glu gene has been functionally transferred to the nuclear genome or the cytosolic tRNA-Glu is now targeted to the plastid and plays a double role in tetrapyrrole and organellar translation in *Lophophytum* and *Ombrophytum*. *Balanophora* spp. maintain an aberrant plastid-encoded tRNA-Glu, which retains key sequence determinants necessary for interactions with the glutamyl-tRNA synthetase (GluRS) and the glutamyl-tRNA reductase (GluTR), suggesting its sole functional role in tetrapyrrole biosynthesis (Su, et al. 2019; Ceriotti, et al. 2025). MSR-seq expression data suggest that the aberrant plastid-encoded tRNA-Glu gene in *B. laxiflora* may be functional because it is expressed and CCA-tailed, although very few reads were obtained (**Table 1**). We did not find a plastid-like tRNA-Glu gene in the transcriptomes or nuclear genomes of *B. subcupularis* or *B. fungosa*. The foreign and mitochondrial-encoded tRNA-Glu gene in *L. pyramidale* was expressed and CCA-tailed, while no mitochondrial tRNA-Glu gene is present in *Balanophora* spp. (**Table 1**).

We evaluated the subcellular localization of enzymes that have the potential to directly interact with tRNA-Glu (GluTR and cytosolic/organellar GluRSs) in *Balanophora* and *Lophophytum* to understand whether they are targeted to the plastids and/or the mitochondria. Specifically, we fused putative transit peptides from these enzymes to GFP and transiently expressed them in *Nicotiana benthamiana* leaves through *Agrobacterium* infiltration. The three key enzymes involved in tetrapyrrole biosynthesis exhibit N-terminal extensions in the three examined species, *L. mirabile*, *L. pyramidale*, and *B. fungosa* (**Dataset 1**). Among these, GluTR shows robust evidence of exclusive plastid targeting. Both *in silico* predictions and microscopy support its plastid localization in all the species (**Table 2**, **Figures 5 and S7**), indicating that the initial steps of tetrapyrrole biosynthesis occur in plastids.

**Figure 5.**
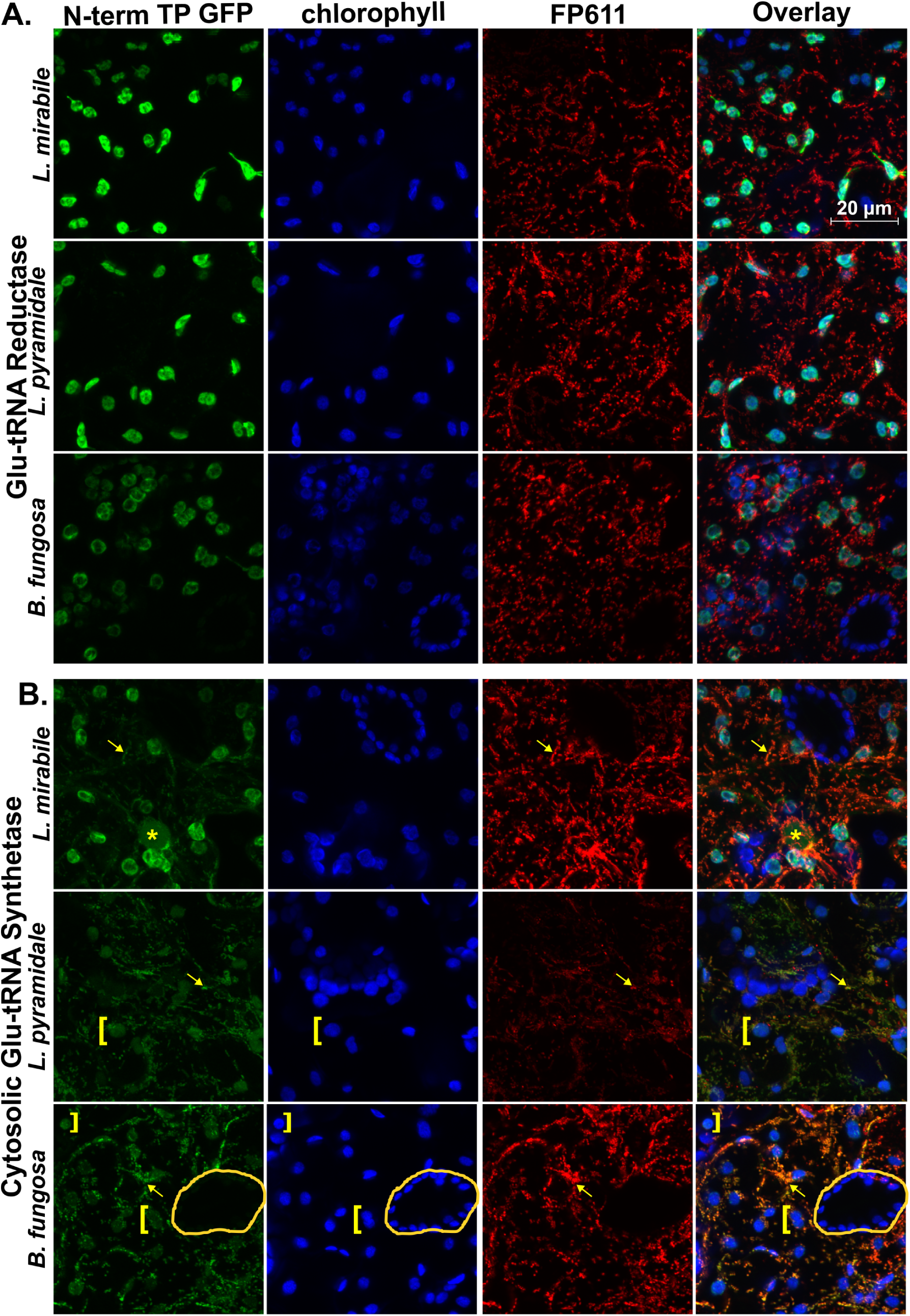
Transit peptides from Balanophoraceae target GluTR and cytosolic GluRS to one or both organelles. Fluorescent confocal images of localization for GFP fused to N-terminal transit peptides from (A) GluTR or (B) cytosolic-type GluRS from three different species. Chlorophyll autofluorescence and IVD-FP611 are used to visualize plastids and mitochondria, respectively. All GluTR transit peptides exhibit strong plastid localization, whereas all cytosolic GluRS transit peptides show evidence of dual localization to mitochondria and plastids (to varying extents). The asterisk denotes nuclear GFP localization of the cytosolic GluRS. Yellow brackets denote plastids exhibiting weak GFP localization. Arrows denote strong mitochondrial colocalization. A pair of untransformed guard cells are circled in the B. fungosa cytosolic GluRS image to demonstrate that weak plastid GFP signal is not due to overlapping bands of chlorophyll autofluorescence. A similar image with untransformed guard cells for L. pyramidale can be seen in Supp. Fig 7C. For negative (no GFP) control, see Supp. Fig 7B.

**Table 2.** Subcellular localization of enzymes that directly interact with tRNA-Glu in *Balanophora fungosa* and *Lophophytum* spp.

| Species | Gene | Organellar Target Prediction |  | Microscopy |
| --- | --- | --- | --- | --- |
|  |  | Localizer | TargetP |  |
| *Balanophora fungosa* | cyt-GluRS | - | Mitochondria | Mitochondria + Plastid (weak) |
|  | org-GluRS | - | - | Undetermined |

|  | GluTR | Plastid | - | Plastid |
| --- | --- | --- | --- | --- |
| *Lophophytum mirabile* | cyt-GluRS | - | Mitochondria | Mitochondria + Plastid + Nucleus (weak) |
|  | org-GluRS | Plastid | Mitochondria | Mitochondria (weak) + Plastid + Nucleus (weak) |
|  | GluTR | Plastid | Plastid | Plastid |
| *Lophophytum pyramidale* | cyt-GluRS | Mitochondria | Mitochondria | Mitochondria + Plastid (weak) |
|  | org-GluRS | Mitochondria | - | Plastid |
|  | GluTR | Plastid | Plastid | Plastid |

The organellar GluRS, typically dual-targeted to plastids and mitochondria in angiosperms, displays a complex localization pattern in Balanophoraceae. *In silico* targeting predictions vary between the three species and between prediction tools. Microscopy revealed plastid localization in *Lophophytum* spp., though weak mitochondrial and nuclear signals were also observed in *L. mirabile*. For the *B. fungosa* enzymes, transient expression assays yielded inconclusive results, leaving its subcellular localization (nuclear, cytosolic, or plastidic) unresolved **(Figure S7)**.

The cytosolic GluRS homolog possesses an N-terminal extension but is truncated in all three species (also in *O. subterraneum* and *Rhopalocnemis phalloides*), missing the first ∼200 aa **(Dataset 1)**. This truncation results in the loss of the GST_C domain (Glutathione S-transferase, C-terminal domain PF00043), a conserved region present across angiosperms. Regarding subcellular localization, *in silico* analyses predict mitochondrial targeting for this truncated protein in the three species. Microscopy results revealed localization to both organelles, although with distinct patterns: mitochondrial and weak plastid signals in *B. fungosa* and *L. pyramidale*, and predominantly plastid with mitochondrial and minor nuclear signals in *L. mirabile*.

### Complete native subunit content and signals of plastid-nuclear coevolution in organellar ribosomes

Given the extreme reductive evolution characterizing Balanophoraceae plastomes, we examined the composition and evolutionary history of their ribosomes. Beyond the high AT bias and elevated substitution rates of the plastid DNA, the reduction in gene content involved both gene loss and IGT to the nucleus. Of the 21 plastid-encoded ribosomal genes present in *Arabidopsis* (Scarpin, et al. 2022), five to seven were transferred to the nucleus in Balanophoraceae, while seven to eleven remain in the plastome (**Figure S8**). The remaining five to seven were not detected in transcriptomic or genomic data, indicating their probable loss in this lineage. In parallel, the mitoribosome of *Lophophytum* spp. is shaped by extraordinary levels of HGT, which has replaced several native mitochondrial ribosomal protein genes (**Figure S8**). Additionally, we identified the putative loss of *rps7*, which was absent from both the mitochondrial and nuclear genome datasets of all Balanophoraceae analyzed.

These genomic perturbations could potentially disrupt cytonuclear coevolution, leading to: (1) ribosome simplification via loss of nuclear-encoded subunits; (2) co-acquisition of foreign nuclear genes to maintain compatibility; or (3) accumulation of compensatory mutations in nuclear genes to restore compatibility. To test these scenarios, we analyzed the ribosomal composition in four holoparasitic species: *L. mirabile*, *L. pyramidale*, *O. subterraneum*, and *B. fungosa*. Using *Arabidopsis* sequences as queries (Waltz et al. 2019), we recovered transcripts corresponding to the full complement of 71 nuclear-encoded mitoribosomal subunits and 33 plastid ribosomal subunits across all four Balanophoraceae (**Figure S8, Tables S7-S8**). These subunits were equally present in the hemiparasitic and autotrophic reference species. Thus, we found no evidence for the loss of nuclear-encoded ribosomal subunits. Furthermore, phylogenetic analyses of all nuclear-encoded organellar subunits revealed a consistent pattern of vertical inheritance. Homologs from Balanophoraceae formed well-supported clades (bootstrap support ≥70%) with other Santalales or grouped within expected lineages (trees available via https://github.com/lfceriotti/Balanophoraceae-MSRseq). Despite the extensive HGT observed in the mitochondrial genomes of *Lophophytum* spp., we detected no evidence of HGT affecting the corresponding nuclear-encoded ribosomal genes.

To evaluate potential compensatory evolution, we assessed synonymous (dS) and non-synonymous (dN) substitution rates. We observed a significant divergence in evolutionary dynamics between nuclear genes targeted to plastids versus those targeted to mitochondria (**Figure S9; Tables S7-S8**). Nuclear genes encoding plastid ribosomal proteins exhibited elevated substitution rates in Balanophoraceae compared to autotrophic relatives, particularly at non-synonymous sites, indicating substantial amino acid divergence (**Figure 6**). However, functional domains remained conserved, and dN/dS ratios were consistently <1 (**Table S8**), indicative of purifying selection retaining essential functionality. This pattern aligns with a scenario of compensatory plastid-nuclear coevolution driven by highly divergent and AT-rich plastid rRNAs and proteins, as well as relaxed selection on plastid translation. In sharp contrast, nuclear genes encoding mitochondrial ribosomal proteins evolved at markedly slower rates, displaying high amino acid sequence conservation (low dN) even in cases of elevated synonymous divergence (dS). In *Lophophytum* spp., this constraint likely facilitates functional compatibility with foreign mitochondrial-encoded subunits acquired from Fabaceae hosts. Collectively, these findings highlight a striking asymmetry: while plastid ribosomes undergo rapid evolution, mitochondrial ribosomes maintain cytonuclear compatibility through extreme sequence conservation despite their chimeric origins.

**Figure 6.**
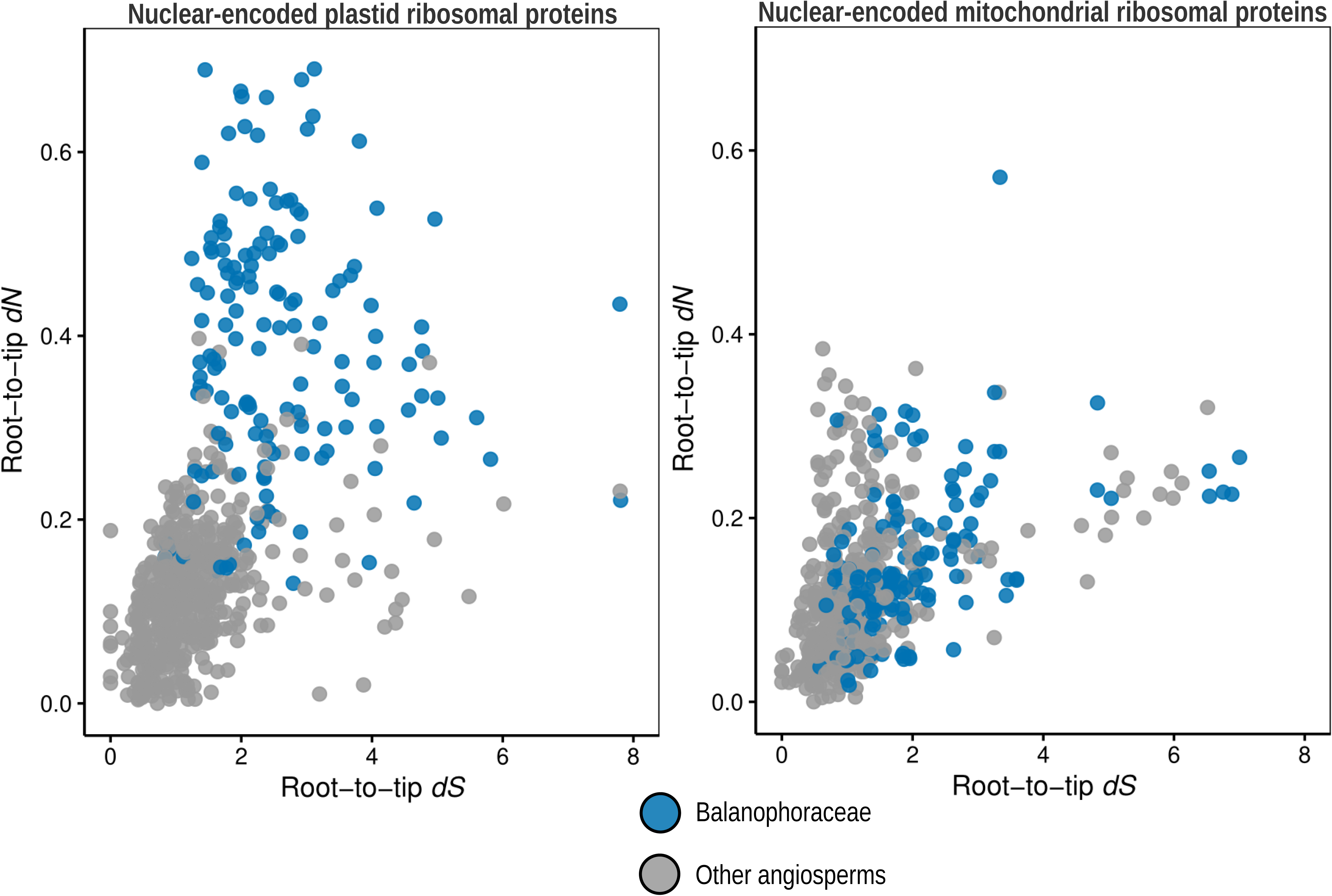
Asymmetric evolutionary rates of nuclear genes encoding organellar ribosomal proteins. Comparison of root-to-tip non-synonymous (dN) versus synonymous (dS) substitution rates for nuclear genes encoding plastid (left panel) and mitochondrial (right panel) ribosomal proteins. Each dot represents a single gene. Blue circles denote orthologs from Balanophoraceae species (*Lophophytum mirabile*, *L. pyramidale*, *Ombrophytum subterraneum*, and *Balanophora fungosa*), whereas grey circles represent orthologs from other autotrophic and hemiparasitic angiosperms included in the reference dataset. Note the distinct elevation in dN values for plastid-targeted genes in Balanophoraceae, indicating accelerated amino acid divergence driven by relaxed purifying selection or compensatory evolution. In contrast, mitochondrial-targeted genes in Balanophoraceae largely overlap with the reference angiosperms, reflecting strong functional conservation despite the holoparasitic lifestyle.

## DISCUSSION

Loss of photosynthesis and a holoparasitic lifestyle leads to a cascade of changes that strongly impact the organelles (Cai 2023). We focused on the organellar translation systems, which require both independent and redundant machinery (e.g. ribosomal subunits) and dual-targeted proteins (e.g. aaRS). In particular, we evaluated the extent of alteration of the translation systems in holoparasites of the family Balanophoraceae (**Figure 7**), some of which have also been heavily impacted by mitochondrial HGT (Sanchez-Puerta, et al. 2017; Roulet, et al. 2020; Roulet, et al. 2024).

**Figure 7.**
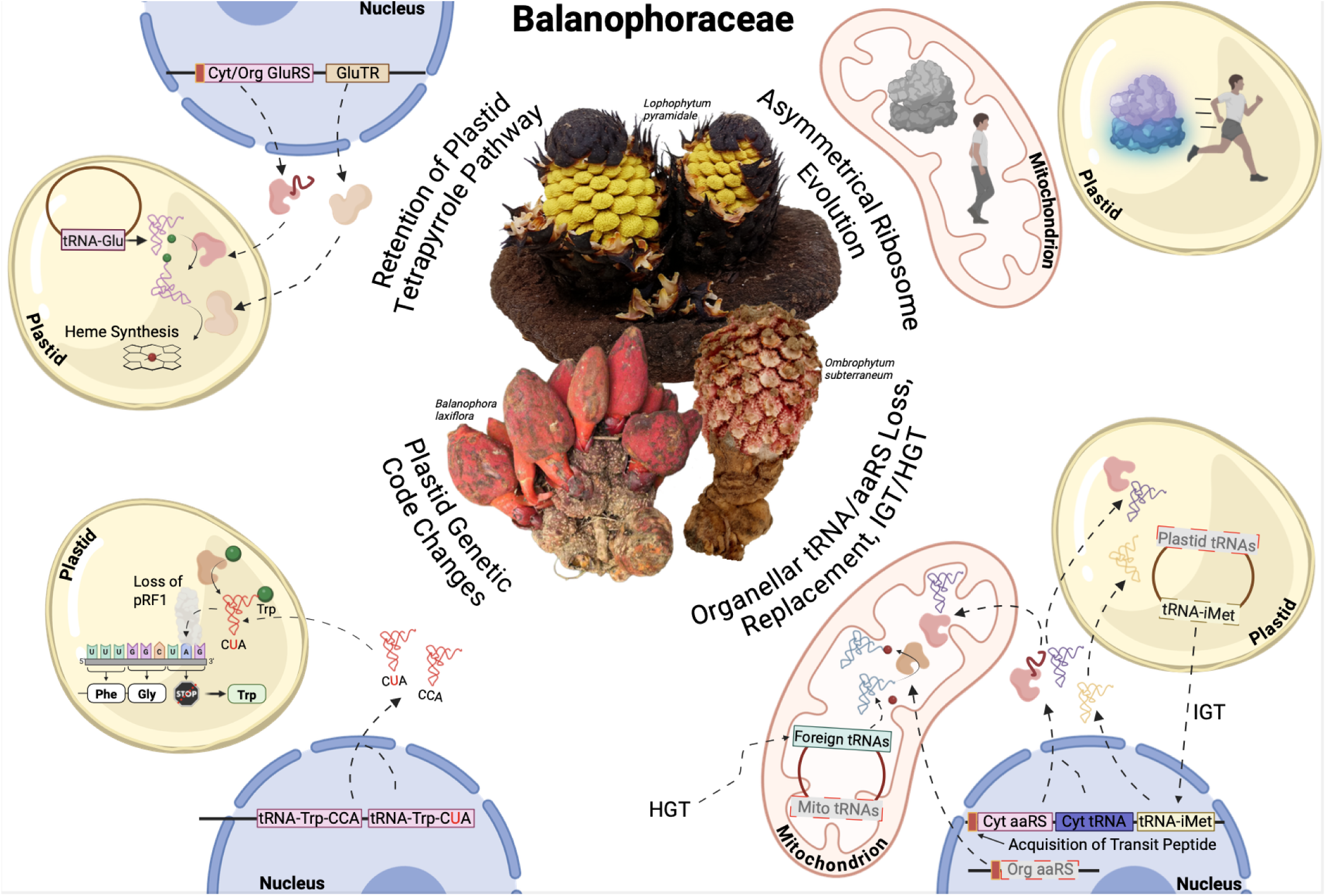
Graphical summary of evolutionary history of organellar translation and tRNA metabolism in the family Balanophoraceae (although note that not all processes described are observed in all species). Boxes drawn with a gray background, gray text, and a dashed border are indicative of extensive loss of that gene class. Figure created with BioRender.

### Evolutionary dynamics of organellar tRNAs: Intracellular transfer and functional integration of foreign genes

The reduction of the plastome in Balanophoraceae involved massive tRNA losses in the common ancestor, leaving the tRNA-Glu gene as the sole survivor in most lineages. While the retention of this gene is evolutionarily expected given its dual role in protein translation and tetrapyrrole biosynthesis (Barbrook, et al. 2006), its preservation is not universal (Bellot and Renner 2016; Klimpert, et al. 2022; Garrett, et al. 2023) with some lineages exhibiting complete plastome loss (Molina, et al. 2014; Cai, et al. 2021; Banerjee and Stefanović 2023; Yu, et al. 2026). In particular, *Lophophytum* and *Ombrophytum* have lost the plastid tRNA-Glu entirely, while *Balanophora* spp. retain an aberrant version (Su, et al. 2019). We found no genomic or transcriptomic evidence of tRNA-Glu gene transfer to the nucleus, and our MSR-seq dataset confirms that the aberrant plastid tRNA-Glu gene in *B. laxiflora* is expressed and CCA-tailed. While read counts are low, other lines of evidence (see below) also support the functionality of the plastid-encoded tRNA-Glu in *Balanophora* spp. in tetrapyrrole biosynthesis.

We identified a striking mechanism of compensation in response to one of these losses: the functional IGT of the plastid initiator tRNA-iMet gene to the nucleus. We confirmed this transfer through expression profiling, which showed high abundance and CCA-tailing of the transcript, and by genomic validation in *B. fungosa* and *B. subcupularis*. This transfer likely occurred in the Balanophoraceae ancestor prior to the loss of the tRNA-iMet gene from the plastome, although the phylogeny of this gene is ambiguous and independent losses and transfers are also possible. Functionally, this nuclear-encoded tRNA-iMet must be recognized by the translation machinery to initiate organellar protein synthesis. We propose a specific pathway: the tRNA is likely charged by a dually targeted cytosolic MetRS (because the organellar MetRS is missing) and subsequently modified by the organellar methionyl-tRNA formyltransferase (MTF), which is conserved in all analyzed species. Additional assays, such as charged tRNA sequencing (Evans, et al. 2017; Watkins, et al. 2022) and organelle purification, are necessary to demonstrate aminoacylation and localization within the plastid compartment. This potentially represents only the second known case of a functional organellar tRNA transfer to the nucleus in eukaryotes, paralleling the recent discovery of plastid tRNA-Pro import in the lycophyte *Selaginella* (Berrissou, et al. 2024).

The mitochondria of Balanophoraceae harbor a chimeric tRNA landscape comprising native genes, plastid-derived tRNAs, and foreign genes acquired via HGT. Expression patterns indicate that the majority of these foreign genes, including a tRNA-Trp located within a foreign MTPT, are transcriptionally active and processed. Conversely, the transcriptional silence of specific loci, such as the foreign tRNA-eMet in *L. pyramidale* and the native tRNA-Glu and tRNA-Ser in *O. subterraneum*, suggests ongoing pseudogenization or that these genes never became functional upon transfer. Notably, the functional integration of foreign tRNAs appears to follow a replacement model, wherein foreign homologs functionally substitute for native genes before or soon after being lost from the mitochondrial genome. This dynamic is strongly corroborated by the strict retention of the cognate organellar aaRSs. While aminoacylation assays are required for definitive proof of charging of these putatively functional tRNAs, current data suggest these foreign transcripts serve as functional substrates for mitochondrial translation. Although MSR-seq can be used to assay aminoacylation (Watkins, et al. 2022), our preliminary investigations found that the issues with RNA extraction from Balanophoraceae tissue made the approaches unreliable (see Data Availability).

### Retention of dual-targeted aaRSs is dictated by organellar tRNA gene content

The loss of organellar tRNA genes is accompanied by the loss of the corresponding nuclear-encoded organellar aaRSs, which are dual-targeted and encoded by nuclear genes (Duchêne, et al. 2005). Previous studies in non-photosynthetic angiosperms suggested that aaRS retention correlates primarily with plastid tRNA content, often uncoupling from mitochondrial requirements (DeTar, et al. 2024). In contrast, our data from Balanophoraceae reveal a stricter coevolutionary scenario where both plastid and mitochondrial tRNA repertoires dictate enzyme retention. We observe a near-perfect correspondence: the presence of a functional organellar tRNA (whether native or foreign) is matched by the retention of its cognate organellar aaRS. Conversely, the loss of specific tRNA isoacceptors matches the loss of the corresponding aaRS, likely due to the relaxation of purifying selection on the enzyme. This correlation validates our expression data; for instance, the retention of the organellar TrpRS in *Lophophytum* spp. and in *O. subterraneum* strongly argues for the functionality of the mitochondrial-encoded tRNA-Trp, which is embedded within a MTPT.

A striking feature of this coevolution is the compatibility between native enzymes and foreign substrates, as small changes in sequence can sometimes create major incompatibilities between mitochondrial tRNAs and aaRSs (Meiklejohn, et al. 2013). In multiple instances, native organellar aaRSs must charge foreign tRNAs. The maintenance of mitochondrial translation in these chimeric systems suggests that the identity elements required for synthetase recognition (Giegé and Eriani 2023) are sufficiently conserved between the parasite and the host-derived tRNAs. As noted regarding ribosomal proteins (Ceriotti, et al. 2021), the high sequence conservation of mitochondrial components across angiosperms likely minimizes cytonuclear incompatibility (Gatica-Soria, et al. 2024). This flexibility implies that organellar aaRSs possess broad substrate specificity, allowing them to accommodate foreign or even cytosolic tRNAs without requiring compensatory evolutionary adjustments. However, it also possible that these aaRSs have evolved to alter their substrate specificity and improve binding/aminoacylation activity with the foreign tRNAs, which could be tested with in vitro aminoacylation assays [e.g., (Gamper and Hou 2020)].

### The plastid genetic code change in *Balanophora* is aided by an anticodon change in a nuclear-encoded tRNA-Trp

The deviations in the plastid genetic code observed in Balanophoraceae, specifically the reassignment of UAG and UGA codons to Trp in *Balanophora* spp. and in *Lophophytum* spp. and *O. subterraneum*, respectively, represent the only documented plastid code changes across angiosperms (Su, et al. 2019; Ceriotti, et al. 2021). This evolutionary shift aligns with the codon capture theory that follows the codon disappearance mechanism (Osawa and Jukes 1989). In line with this theory, the extensive reduction of these plastomes coupled with a strong AT mutational bias led to the disappearance of UGA or UAG codons from the few remaining protein-coding genes (Su, et al. 2019; Ceriotti, et al. 2021), creating a permissive environment for the loss of the canonical peptide RFs responsible for their recognition. Indeed, we confirmed the loss of pRF1 in *Balanophora* spp. and pRF2 in *Lophophytum* spp. and *O. subterraneum*, a prerequisite that allowed these former stop codons to become translatable.

While the reassignment of UGA to Trp is a frequent event in mitochondrial genomes, the reassignment of UAG to Trp is exceptionally rare (Sengupta, et al. 2007). We propose that in *Balanophora*, this transition is mechanistically driven by the import of a duplicated nuclear tRNA-Trp with a mutated CUA anticodon, which we identified in the nuclear genomes of *B. fungosa* and *B. subcupularis*. This tRNA is likely targeted to the plastid, enabling the translation of UAG as Trp by the cytosolic TrpRS, given that the organellar TrpRS is missing, and facilitated by the absence of pRF1. A change in the CCA anticodon is necessary because cognate tRNA-Trp cannot structurally recognize UAG (Heaphy, et al. 2016). A critical question arises regarding the potential interference of this tRNA with cytosolic translation, where UAG functions as a canonical stop codon. We hypothesize that the high efficiency and abundance of cytosolic RFs prevent the misuse of this suppressor tRNA in the cytoplasm, a competition dynamic well-documented in systems where RF depletion promotes stop codon readthrough (Korkmaz, et al. 2014). Conversely, in the plastid, the verified absence of pRF2 in *Lophophytum* spp. and *O. subterraneum* likely facilitates the translation of UGA by the canonical tRNA-Trp-CCA. Although typically restricted to UGG, biochemical evidence demonstrates that this tRNA acts as a preferential near-cognate suppressor at UGA codons. This misreading event is strongly favored in the absence of kinetic competition from pRF2 (Freistroffer, et al. 2000; Sengupta, et al. 2007; Korkmaz, et al. 2014).

### Reconfiguration of tRNA-Glu interactions in Balanophoraceae plastids

Within Balanophoraceae, the evolution of the tRNA-Glu gene spans the broadest range of scenarios known for any eukaryotic lineage with primary plastids. Some genera, such as *Sarcophyte* and *Thonningia*, appear to retain the canonical dual function of the gene, in *Balanophora* the tRNA-Glu gene is predicted to be functional solely for tetrapyrrole synthesis, and in *Lophophytum* and *Ombrophytum* the gene has been entirely lost (Su, et al. 2019; Ceriotti, et al. 2021; Kim, et al. 2023; Ceriotti, et al. 2025). The striking variability displayed by the tRNA-Glu gene sequence in Balanophoraceae highlights the degree of divergence the gene can tolerate once selection pressure on plastid protein synthesis has been alleviated. Despite this variability, several signatures of purifying selection remain. When present, the tRNA-Glu gene consistently occupies a conserved genomic position and exhibits a GC content ∼2-fold higher than the plastome average (Su, et al. 2019; Ceriotti, et al. 2025). While highly aberrant, the tRNA-Glu is present in the plastome of all *Balanophora* species examined (Svetlikova, et al.; Chen, et al. 2019; Su, et al. 2019). Across the family, the gene retains key sequence determinants required for interaction with the organellar GluRS and GluTR, supporting its continued role in tetrapyrrole biosynthesis (Su, et al. 2019; Ceriotti, et al. 2021; Ceriotti, et al. 2025). By contrast, most polymorphisms are concentrated in the D-loop and anticodon stem–loop, producing noncanonical or entirely absent anticodons and likely precluding its function in translation (Su, et al. 2019; Ceriotti, et al. 2021; Ceriotti, et al. 2025). In these species, the plastid translational function of the tRNA-Glu gene is presumably replaced by the import of the cytosolic counterpart. In lineages where the plastid gene has been lost, the full functional replacement by the cytosolic tRNA-Glu has been proposed based on the presence of sequence determinants in this nuclear gene (Ceriotti, et al. 2021).

To deepen our understanding of plastid tRNA-Glu interactions, we examined the subcellular localization of enzymes that directly interact with this tRNA in *Lophophytum* spp., which lack the plastid tRNA-Glu gene, and in *Balanophora*, where the tRNA-Glu gene is extremely divergent. Most of the transit peptides tested in our *in vivo* microscopy assays localized to mitochondria and/or plastids; however, the specific organelle sometimes differed from one or both *in silico* predictions. The heterogeneity observed among *Lophophytum* species was particularly unexpected given their shared evolutionary scenario regarding the plastid tRNA-Glu gene.

Several factors help explain the challenges of predicting, or experimentally assessing, protein targeting in heterotrophic plants (DeTar, et al. 2024). Beyond the inherent limitations of targeting prediction algorithms (Sperschneider, et al. 2017; Almagro Armenteros, et al. 2019), predictive accuracy in taxa that remain uncharacterized is compromised by the extensive diversity in both transit peptide sequence composition and protein import machinery among plant lineages (Christian, et al. 2020). Moreover, some organellar proteins are known to be imported without N-terminal transit peptides (Chen, et al. 2004; Neupert and Herrmann 2007). These issues are exacerbated by pronounced divergence in the plastid import machinery of many heterotrophic plants (Guo, et al. 2023; DeTar, et al. 2024), which further reduces the sensitivity of models trained mainly on photosynthetic species. Heterologous expression assays also introduce artifacts, as GFP-fusion localization can be influenced by protein folding efficiency, expression levels, leaf developmental stage, and the truncation of internal targeting sequences (Tanz, et al. 2013; Sharma, et al. 2018; Jeong, et al. 2022). Finally, incomplete or uncertain protein models, particularly in the absence of long-read RNA sequencing, may obscure true N-terminal regions, preventing the detection of alternative splicing or alternative transcription/translation initiation events that generate differentially targeted isoforms.

Despite these difficulties, our results unambiguously support plastid targeting of GluTR. Both *in silico* analyses and *in vivo* localization experiments indicate that the first committed step of tetrapyrrole biosynthesis occurs in this organelle regardless of the presence or absence of the plastid tRNA-Glu gene. The adaptation to a different substrate, the imported cytosolic tRNA-Glu in *Lophophytum* spp. and an extremely divergent tRNA-Glu in *Balanophora*, point to a diversification in substrate recognition for GluTR across Balanophoraceae, a diversification likely to be even more extensive given the tRNA-Glu gene divergence in the family (Ceriotti, et al. 2025). In addition, our results support the acquisition of an N-terminal transit peptide to the organelles in the cytosolic GluRS. The identical position of this transit peptide and the concurrent loss of the first ∼200 amino acids in all three species examined suggest an ancestral acquisition of organelle targeting.

Because interactions between organellar and cytosolic aaRSs and tRNAs could generate cytonuclear incompatibilities, we propose the following hypothesis as the most consistent with our observations. The cytosolic GluRS in *Lophophytum* would be responsible for charging the imported cytosolic tRNA-Glu within plastids, where it would support both tetrapyrrole biosynthesis and translation. In *Balanophora*, this enzyme may support translation in plastids and mitochondria. The organellar GluRS, in turn, might charge the mitochondrial and the plastid tRNA-Glu in *Lophophytum* and *Balanophora*, respectively, as these tRNAs are expressed and post-transcriptionally modified and therefore likely functional. Unfortunately, our current results cannot conclusively determine which organelle(s) receive the cytosolic and organellar GluRS proteins, nor the aminoacylation and functionality of the tRNAs. Also, redundant tRNA import pathways may operate before mt-tRNA loss in the parasites, as has been shown in *Silene* (Warren, et al. 2021). Future work integrating organellar tRNA-import analyses, *in vitro* aminoacylation assays, and comparative proteomics will be crucial for resolving the evolution of this complex cytonuclear system in Balanophoraceae.

### Cytonuclear compatibility and the asymmetry of organellar coevolution

Another intriguing aspect of the organellar translation under extreme heterotrophic conditions is the function of the organellar ribosomes, which rely on coordinated expression of subunits encoded by both the nuclear and organellar genomes, a process that requires cytonuclear coevolution (Burton and Barreto 2012; Sloan, et al. 2022). Disruption of this coevolution can impair organellar ribosomal function and compromise organismal fitness (Wolff, et al. 1994; Latorre-Pellicer, et al. 2016; Sloan, et al. 2018). Although the Balanophoraceae exhibit the full complement of nuclear-encoded subunits that form part of the mitochondrial and plastid ribosomes, cytonuclear coevolution may be affected in two major ways. First, plastid ribosomes were impacted by the loss of photosynthesis, which was followed by accelerated substitution rates, loss of ribosomal genes, and extreme AT nucleotide bias (>85%) (Schelkunov, et al. 2019; Su, et al. 2019; Ceriotti, et al. 2025). Indeed, we observed a correlated acceleration in the evolutionary rates of interacting nuclear genes encoding ribosomal subunits in the Balanophoraceae. While such correlated evolution is frequently cited as a hallmark of compensatory coevolution in lineages with divergent organellar genomes (Sloan, et al. 2013; Zhang, et al. 2015; Rockenbach, et al. 2016; Weng, et al. 2016; Tressel, et al. 2025), relaxation of purifying selection represents an equally important alternative hypothesis. Given that the plastid no longer supports the heavy translational demand of the photosynthetic apparatus (Raven 1995; Li, et al. 2017; Heinemann, et al. 2020), the functional constraints on plastid ribosomes are likely reduced. Consequently, the accelerated rates in nuclear subunits may not solely reflect adaptive compensation to restore optimal function, but rather the accumulation of slightly deleterious mutations permitted by the reduced selective pressure on the plastid translational machinery.

Second, mitochondrial ribosomes in *Lophophytum* spp., though typically evolving at slower rates, may be influenced by the frequent acquisition of foreign DNA replacing several of the native ribosomal proteins with foreign homologs in the mitochondrial genome (Sanchez-Puerta, et al. 2017; Roulet, et al. 2024). This contrasts sharply with *Ombrophytum* and *Balanophora*, which retain native mitochondrial genes, providing a robust comparative framework within the family. Given the massive mitochondrial HGT in *Lophophytum* spp. involving donors phylogenetically distant from the parasitic plants, we anticipated signs of mitonuclear incompatibility or, alternatively, rapid compensatory evolution in the nuclear-encoded subunits. Unexpectedly, our results contradict this prediction. Nuclear genes encoding mitochondrial proteins in *Lophophytum* spp. display high sequence conservation and low non-synonymous substitution rates, showing no evidence of the accelerated evolution as observed in the plastid system. This suggests that the foreign mitochondrial genes possess a remarkable capacity for functional integration without necessitating compensatory changes in the nuclear background. These observations align well with similar findings when analyzing the chimeric OXPHOS complexes in *Lophophytum* spp., which exhibit a complete set of subunits (Gatica-Soria, et al. 2022) and no evidence of compensatory evolution (Ceriotti, et al. 2022), while the respiratory function was similar to that of free-living plants (Gatica-Soria, et al. 2024).

Collectively, these findings highlight a striking evolutionary asymmetry in the organellar translation machineries of Balanophoraceae. While plastid ribosomes show clear signatures of accelerated evolution driven by a highly mutated and reduced plastome, mitochondrial ribosomes appear to have maintained functionality through the structural conservation of their components. This conservation may have allowed the mitochondrial system to tolerate massive horizontal gene replacement without requiring compensatory nuclear evolution to restore cytonuclear compatibility.

## MATERIALS AND METHODS

### Plant material and RNA extraction

We studied three species of Balanophoraceae and *Arabidopsis thaliana* Col-0. Inflorescences were collected from *Lophophytum pyramidale* in San Ignacio, Misiones, Argentina (individual #3), from *Ombrophytum subterraneum* in Rodeo, Jujuy, Argentina (individual #1), and from two samples of *Balanophora laxiflora* (XHSG and DHSG) on Dayaoshan Mountain, Guangxi province in South China. *Arabidopsis thaliana* Col-0 seeds were surface sterilized for 10 min in a 50% bleach solution with Tween 20 (VWR 0777A) followed by three washes with sterile dH_2_O. Seeds were sown onto MS-agar plates and grown for 3 weeks. Flash-frozen tissues from the inflorescences of the three holoparasites and from rosette leaves of *Arabidopsis* were ground and kept at −80 °C.

For *L. pyramidale*, *O. subterraneum*, and *A. thaliana*, total RNA was extracted using a CTAB protocol modified for highly viscous samples rich in polysaccharides (Zeng and Yang 2012), with modifications such as lower temperature and pH 7.5 to keep the amino acids attached **(Supplementary Note 1)**. For the two individuals of *Balanophora laxiflora*, total RNA was extracted following Yang et al. (Yang, et al. 2008).

### MSR-seq library construction, sequencing, and analysis

Sequencing of tRNAs was performed using an MSR-seq protocol (Watkins, et al. 2022), as described previously (Ceriotti, et al. 2024). Two technical replicate libraries (independent PCR amplifications of the same library bead prep) were generated for the *A. thaliana* Col-0, *L. pyramidale*, and *O. subterraneum* samples. An input of approximately 500 ng of RNA was used except for the *L. pyramidale* libraries, which were made from approximately 300 ng due to lower concentration of that sample. MSR-seq can be used to infer the proportion of tRNAs that are aminoacylated when samples are first treated with sodium periodate followed by sodium tetraborate (Watkins, et al. 2022). For these three species, libraries were generated for both treated and untreated (control) portions of each sample, as described previously (Ceriotti, et al. 2024). However, our initial analyses indicated that the RNA extraction methods necessary for Balanophoraceae tissues did not adequately preserve the aminoacylation state, and treated samples were not analyzed further. All presented results are based on the untreated control libraries only. In a separate round of library construction and sequencing, three technical replicate libraries (untreated only) were generated for each of the two *B. laxiflora* samples. Two of these replicates were from independent PCR amplifications of the same library bead prep, whereas the third was amplified from a separate library bead prep from the same RNA sample. Approximately 300 ng and 200 ng of input RNA was used for the *B. laxiflora* DHSG and XHSG preps, respectively. Library sequencing was performed by Novogene on an Illumina NovaSeq X platform (2×150 bp). The *L. pyramidale*, *O. subterraneum*, and *A. thaliana* libraries were sequenced together in one run, and the *B. laxiflora* libraries were sequenced together in a later run.

MSR-seq reads were mapped to a tRNA reference database to quantify read abundance and detect post-transcriptional modifications, including additions of CCA tails. This analysis was performed with a previously published pipeline (Ceriotti, et al. 2024). Briefly, BBMerge (Bushnell, et al. 2017) was used to combine each pair of R1 and R2 reads into a single sequence, which were then further trimmed with a custom script to remove the adapter sequences from the MSR-seq CHO (capture hairpin oligo). Read pairs that could not be combined by BBMerge or did not have the expected adapter sequence were excluded from downstream steps. All successfully processed reads were mapped against a reference database (**Dataset 2**) containing all annotated organellar tRNAs from *A. thaliana*, *B. laxiflora*, *L. pyramidale*, and *O. subterraneum*. It also included all nuclear tRNAs from *A. thaliana* from the plantRNA 2.0 database (Cognat, et al. 2022). After an initial mapping round, we discover a plastid-like nuclear tRNA-iMet in *O. subterraneum* MSR-seq data, which was included in subsequent mapping rounds. Comprehensive annotations of Balanophoraceae nuclear tRNAs are not available, but we did add specific nuclear tRNAs of interest to the reference database. Specifically, we included nuclear tRNA-Trp-CCA and tRNA-Trp-CUA sequences from nuclear assemblies of *B. yakushimensis* (Yu, et al. 2025) and plastid-like nuclear copies of tRNA-iMet from *B. fungosa* and *B. subcupularis* (Chen, et al. 2023). Finally, the reference database included the sequence of a synthetic spike-in tRNA that was added to each sample as a control (Ceriotti, et al. 2024).

Mapping was conducted with Bowtie v2.2.5 (Langmead and Salzberg 2012), using the following sensitive parameters to facilitate detection of reads with mismatches and indels that were introduced due to post-transcriptional base modifications: -p 12 -L 10 -i C,1 --mp 5,2 --score-min L,-0.7,-0.7. We used a custom local alignment pipeline based on NCBI BLASTN 2.14.1+ (Camacho, et al. 2009) to pre-process reads and trim 5ʹ extensions prior to Bowtie mapping. A custom script was then used to parse the Sequence Alignment Map (SAM) file generated by Bowtie, quantifying the number of reads that mapped to each reference sequence and determining whether they contained a full CCA tail or were lacking one or more 3ʹ nucleotides. SAM files were also parsed to identify reads with mismatches or small indels relative to the reference sequence, which can be caused by reverse transcriptase misincorporations in response to post-transcriptionally modified bases in the RNA template, using a previously published variant calling workflow (Edera and Sanchez-Puerta 2021). We prepared a custom script to extract reads from SAM files that mapped to tRNAs of interest and exhibit specific anticodon modifications (available via https://github.com/lfceriotti/Balanophoraceae-MSRseq). Raw read abundances should be interpreted with caution since they are prone to strong biases in the representation of particular genes. These biases are likely linked to secondary structure, post-transcriptional modifications, and other methodological limitations inherent to tRNA-seq (Ma, et al. 2021; Warren, et al. 2021). For this reason, relative comparisons among samples generally provide the most meaningful insights from MSR-seq data.

### Homology searches

To search for nuclear homologs of aaRSs, organellar RFs, GluTR, and organellar ribosomal proteins, we conducted tBLASTn searches (e-value < 0.001) against the transcriptomes of *Lophophytum mirabile* (Garcia, et al. 2021), *L. pyramidale* (Roulet, et al. 2024), and *O. subterraneum* (Garcia and Sanchez-Puerta 2024), and *Balanophora fungosa* (Schelkunov, et al. 2021), as well as the nuclear genomes of *B. fungosa* and *B. subcupularis* (Chen, et al. 2023). For the ribosomal proteins, we also searched the transcriptomes of three hemiparasitic species within the order Santalales—*Daenikera* sp., *Dendropemon caribaeus*, and *Malania oleifera* (Schelkunov, et al. 2021) and the transcriptome of the photosynthetic host of *L. mirabile*, *Anadenanthera colubrina* (PRJNA1197971). In all cases, we used *Arabidopsis thaliana* protein sequences as queries. Candidate hits with both identity and coverage exceeding 20% were retained for further analysis and verified if they corresponded to plant homologs of the *Arabidopsis* gene used as the original query by reciprocal BLASTp searches. The accession numbers for the cytosolic and organellar aaRSs **(Table S6)** were taken from DeTar *et al*. (2024) and those of GluTR and RFs from Ceriotti *et al*. (2021). Nuclear genes (**Table S9**) involved in mitochondrial ribosomes were retrieved from Waltz *et al*. (2019), and those related to plastid ribosomes from Scarpin *et al*. (2022). Conserved protein domains for ribosomal proteins were annotated via the NCBI Conserved Domain Database (CDD).

### Subcellular localization through GFP assays

We identified homologs of the three enzymes of interest: cytosolic GluRS, organellar GluRS, and GluTR, as described above. The transit peptides that guide protein subcellular localization were predicted using TargetP 2.0 (Almagro Armenteros, et al. 2019) and LOCALIZER (Sperschneider, et al. 2017) software, and multiple sequence alignments including several angiosperms.

The nine transit peptides, along with the subsequent 30 bp of each gene, were synthesized after codon optimization for *Nicotiana benthamiana*, and then used to generate constructs using Gateway cloning technology in *Escherichia coli* strain DH5α as described previously (DeTar, et al. 2024). The vector carried both the mitochondrial control IVD-FP611 and a GFP linked to the C-terminus of the transit peptide of interest. We validated the construct sequences through whole-plasmid sequencing (Plasmidsaurus) and then transformed them into *Agrobacterium tumefaciens* strain C58C1 through electroporation. The nine constructs were transiently transformed into *N. benthamiana* leaves via *Agrobacterium* infiltration and the imaging was performed with a Nikon A1-NiE confocal microscope equipped with a CFI Plan Apo VC 60 XC WI objective. We assessed the subcellular localization of the nine transit peptides based on colocalization of GFP with eqFP611 (mitochondria) or with autofluorescence (chloroplasts) using NIS element viewer (Nikon) software. For four constructs, the infiltration and microscopy assays were repeated. Constructs are available from Addgene (ID numbers 228382-228390).

### Phylogenetic analyses

To determine the phylogenetic origin of nuclear genes encoding organellar ribosomal proteins and aaRSs, orthologous sequences were retrieved from 11 selected photosynthetic species using the PLAZA v5.0 database (Van Bel, et al. 2021) (**Table S10**). Alignments were generated using MAFFT v7.407 (Katoh and Standley 2013), followed by conversion to codon alignments with PAL2NAL (Suyama, et al. 2006). Poorly aligned or highly divergent regions were filtered using BMGE v1.12 (Criscuolo and Gribaldo 2010). Phylogenetic trees were inferred under a maximum likelihood framework using RAxML v8.2.11 (Stamatakis 2014), with the GTR+GAMMA+I substitution model and 1,000 bootstrap replicates. Tree visualization was performed using FigTree (http://tree.bio.ed.ac.uk/software/figtree/). Alignments and trees were deposited in https://github.com/lfceriotti/Balanophoraceae-MSRseq.

### Estimation of evolutionary rates and selective pressure of ribosomal protein genes

To explore potential compensatory evolution, we estimated substitution rates in nuclear genes encoding organellar ribosomal subunits. Analyses were performed using the *codeml* program from PAML v4.7 (Yang 2007), employing the F3×4 codon frequency model. We allowed ω (dN/dS) to vary among branches (model = 1), with initial values set to ω = 0.5 and κ = 2 for plastid genes. The input phylogenies used in PAML analyses were based on the APG IV framework (Group, et al. 2016) and phylogenetic relationships within Santalales (Nickrent 2020). In addition, root-to-tip substitution rates were calculated using the *castor* package in R, using the last common ancestor of eudicots as the root. For each nuclear gene, the dN/dS ratio was computed for terminal branches, serving as a proxy for lineage-specific selection pressure. These analyses were conducted for all nuclear-encoded plastid ribosomal genes, as well as for a representative subset of 35 mitochondrial ribosomal genes.

## Supporting information

Supplemental Figuures

## DATA AVAILABILITY

Raw sequencing reads are available via the NCBI Short Read Archive (SRA) under BioProject PRJNA1393114. Processed data files and code are available via GitHub (https://github.com/lfceriotti/Balanophoraceae-MSRseq).

## ACKNOWLEDGEMENTS

We thank AK Broz for laboratory assistance and ME Roulet and VC Gómez Villafañe for sample collection. This work was supported by funding from the National Science Foundation (MCB-1933590, MCB-2322154, and IOS-2208908), the University of Nebraska Foundation, an HHMI Hanna H. Gray Fellowship, and an IUBMB Wood-Whelan Research Fellowship.

## Supplementary Material

**Table S1.** Summary of MSR-seq libraries generated for this study, including number of reads produced, successfully processed (removal of expected adapter sequencing and merging R1 and R2 reads with BBMerge), and mapped to the tRNA reference set.

**Table S2.** Proportion of processed reads from each species mapping to tRNAs of different cellular compartments in the reference set.

**Table S3.** Processed reads mapping to *Arabidopsis* plastid tRNAs.

**Table S4.** Total number of reads and number of reads ending with CCA that mapped to selected tRNA sequences.

**Table S5.** Number of processed reads mapping to *Arabidopsis* nuclear tRNA-Trp genes. The nuclear-Ath-TrpCCA-183740 gene exhibited much lower coverage than the other nuclear tRNA-Trp genes and was not included in this analysis.

**Table S6.** Aminoacyl-tRNA synthetases (aaRSs) identified in the transcriptome or nuclear genome of Balanophoraceae.

**Table S7.** Estimates of non-synonymous (dN) and synonymous (dS) substitution rates (ω = dN/dS) on terminal branches and root-to-tip paths for nuclear genes involved in mitochondrial ribosomes.

**Table S8.** Estimates of non-synonymous (dN) and synonymous (dS) substitution rates (ω = dN/dS) on terminal branches and root-to-tip paths for nuclear genes involved in plastid ribosomes.

**Table S9.** Nuclear genes encoding organellar ribosome subunits used to search for homologs in the analyzed transcriptomes and genomes of other angiosperms.

**Table S10.** Transcriptomes and genomes used for evolutionary analyses of organellar ribosome protein-encoding genes.

**Figure S1.** CCA tail integrity proportions in the MSR-seq libraries (see Table S1 for library details).

**Figure S2.** Number of reads (read count and parts per million -PPM- of raw sequenced reads) in the MSR-seq libraries (see Table S1 for library details) mapping with best score to *Arabidopsis* plastid tRNA genes. Replicates 1 and 2 were averaged for the *Arabidopsis* and no-template libraries. *Ombrophytum subterraneum* reads mapping to the *Arabidopsis* plastid tRNA-iMet gene are indicated with dashed ellipses.

**Figure S3.** Read coverage and sequence variants in mitochondrial tRNAs. For each species, raw read counts were summed across libraries after excluding reads that are truncated at the 3′ end (lacking >7 nt). Only genes with >50 reads per species are shown.

**Figure S4.** Read coverage and sequence variants in nuclear tRNA-Trp genes in *Balanophora laxiflora* individuals. For each individual, raw read counts were summed across libraries after excluding reads that are truncated at the 3′ end (lacking >7 nt). Some variants, particularly those approaching 100% read coverage, may reflect differences between the genomic DNA sequences of *B. laxiflora* and *B. yakushimensis* rather than post-transcriptional modifications.

**Figure S5.** Read coverage and sequence variants in nuclear tRNA-Trp genes in *Lophophytum pyramidale* and *Ombrophytum subterraneum*. Read counts were summed across replicates for each species after excluding reads that are truncated at the 3′ end (lacking >7 nt). Some variants, particularly those approaching 100% read coverage, may reflect differences between the genomic DNA sequences of *Arabidopsis* and the parasitic plants rather than post-transcriptional modifications.

**Figure S6.** Maximum likelihood phylogenetic trees of aminoacyl-tRNA synthetases (aaRSs) from Balanophoraceae. **A.** Trees providing evidence of foreign aaRSs in Balanophoraceae (in red font). **B.** Trees showing the native origin of aaRSs that interact with functional foreign organellar-encoded tRNAs. Bootstrap support values >50% are shown above the branches. Branch lengths indicate substitutions per site.

**Figure S7.** Transit peptides from *Balanophoraceae* target organellar GluRS to plastids. **A.** Fluorescent confocal images of localization for GFP fused to N-terminal transit peptides from organellar-type GluRS from three different species. Chlorophyll autofluorescence and IVD-FP611 are used to visualize plastids and mitochondria, respectively. **B.** No-GFP control (i.e. just IVD-FP611) visualized under the same settings and conditions as test constructs. **C.** Additional images of localization for GFP construct with an N-terminal transit peptide from cytosolic-type GluRS from *L. pyramidale*. Yellow brackets denote plastids exhibiting weak GFP localization. Note the lack of GFP signal for untransformed guard cell and pavement cell (outlined in yellow) compared to plastid from neighboring transformed cell (yellow bracket) showing true GFP fluorescence.

**Figure S8.** Subunit composition of plastid and mitochondrial ribosomes in Balanophoraceae and *Arabidopsis*, and identification of mitochondrial, plastid or nuclear-encoded genes.

**Figure S9.** Evolutionary dynamics of organellar ribosomes in Balanophoraceae. Maximum likelihood phylogenies of two representative nuclear genes (*rps1* and *rpl23*) encoding plastid-targeted and mitochondrion-targeted ribosomal proteins, respectively. Sequences from Balanophoraceae are highlighted in blue. Conserved protein domains identified in *Balanophora fungosa* are indicated.

