## Supplemental Figuures for "Reshaping Organellar Translation and tRNA Metabolism: The Consequences of Photosynthesis Loss and Massive Horizontal Gene Transfer"

**Figure S1**.CCA tail integrity proportions in the MSR-seq libraries (see Table S1 for library details).

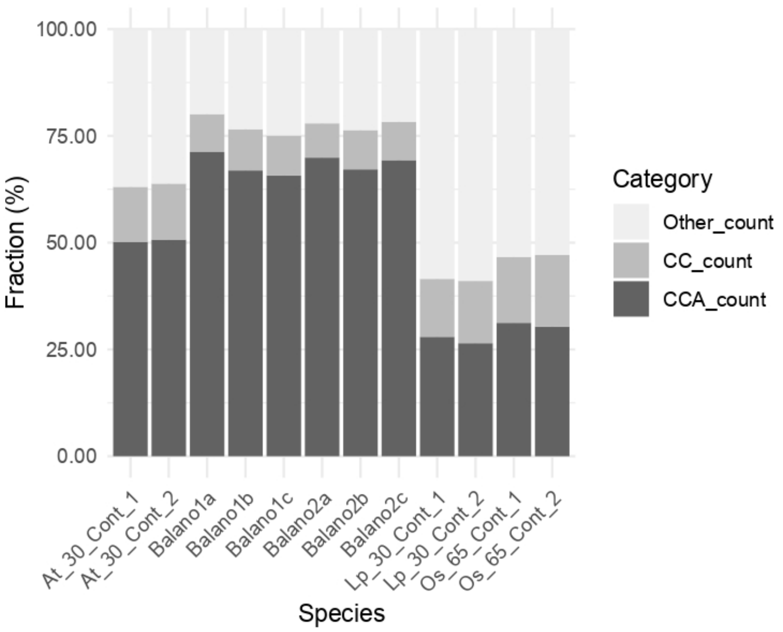

**Figure S2**.Number of reads (read count and parts per million -PPM- of raw sequenced reads) in the MSR-seq libraries (see Table S1 for library details) mapping with best score to *Arabidopsis* plastid tRNA genes. Replicates 1 and 2 were averaged for the *Arabidopsis* and no-template libraries. *Ombrophytum subterraneum* reads mapping to the Arabidopsis plastid tRNA-iMet gene are indicated with dashed ellipses.

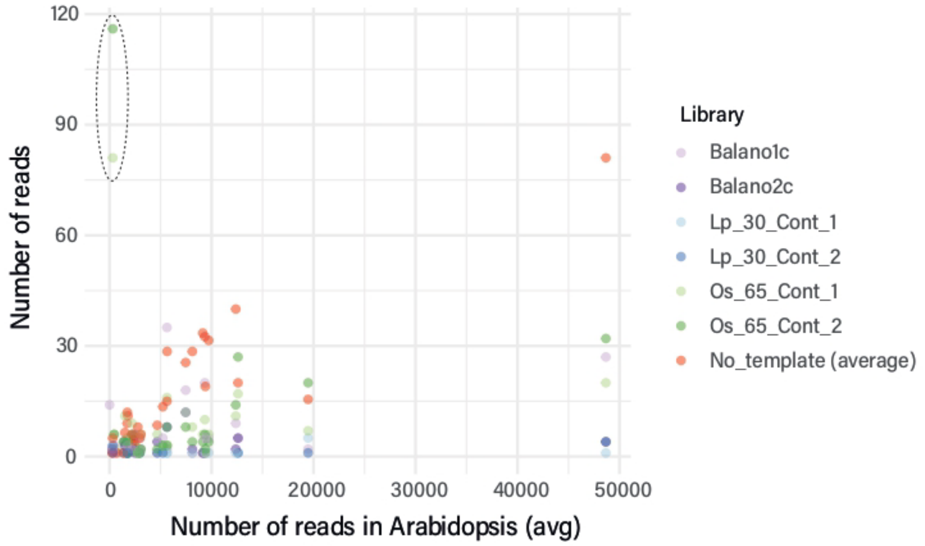


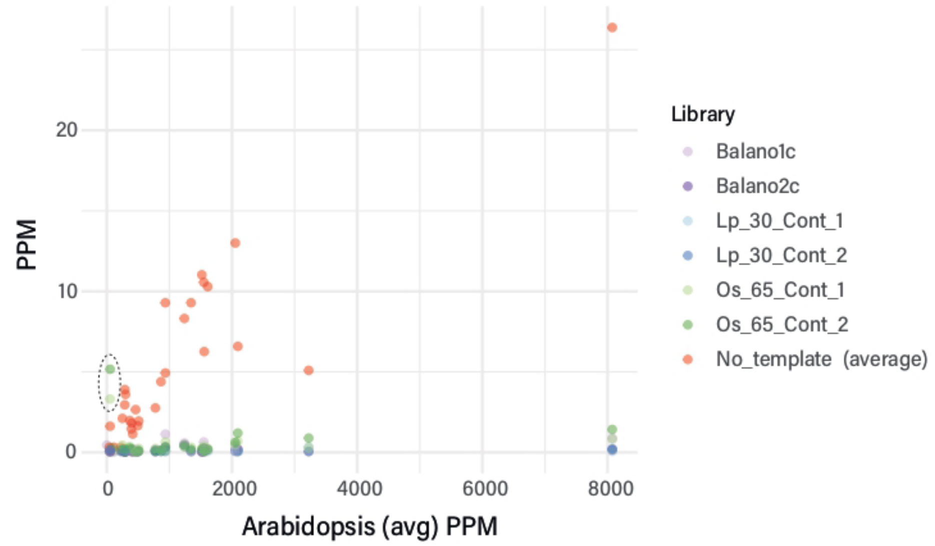

Figure S3. Read coverage and sequence variants in mitochondrial tRNAs. For each species, raw read counts were summed across libraries after excluding reads that are truncated at the 3′ end (lacking >7 nt). Only genes with >50 reads per species are shown.

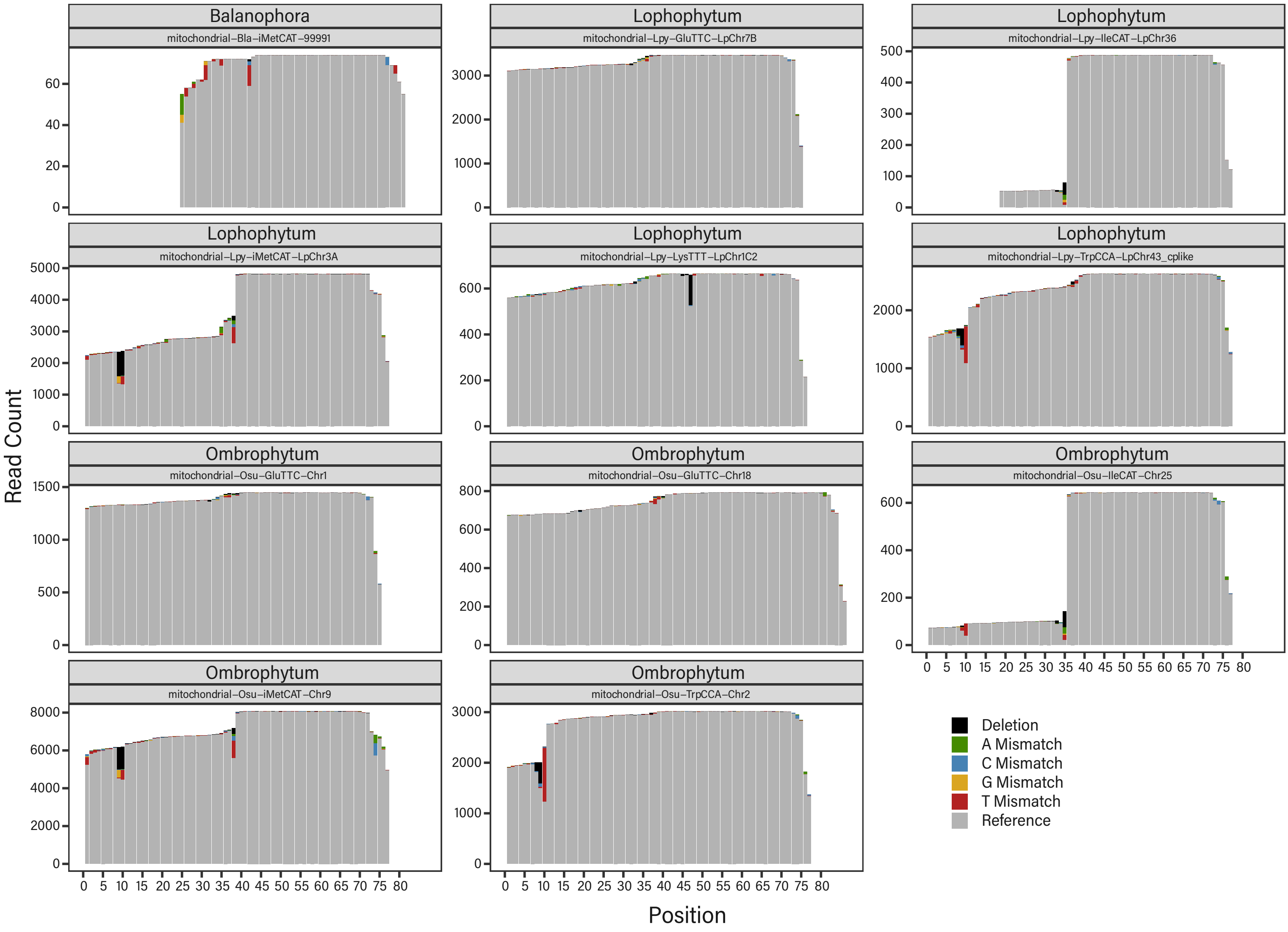

Figure S4. Read coverage and sequence variants in nuclear Trp tRNAs in *Balanophora laxiflora* individuals. For each individual, raw read counts were summed across libraries after excluding reads that are truncated at the 3′ end (lacking >7 nt). Some variants, particularly those approaching 100% read coverage, may reflect differences between the genomic DNA sequences of *B. laxiflora* and *B. yakushimensis* rather than post-transcriptional modifications.

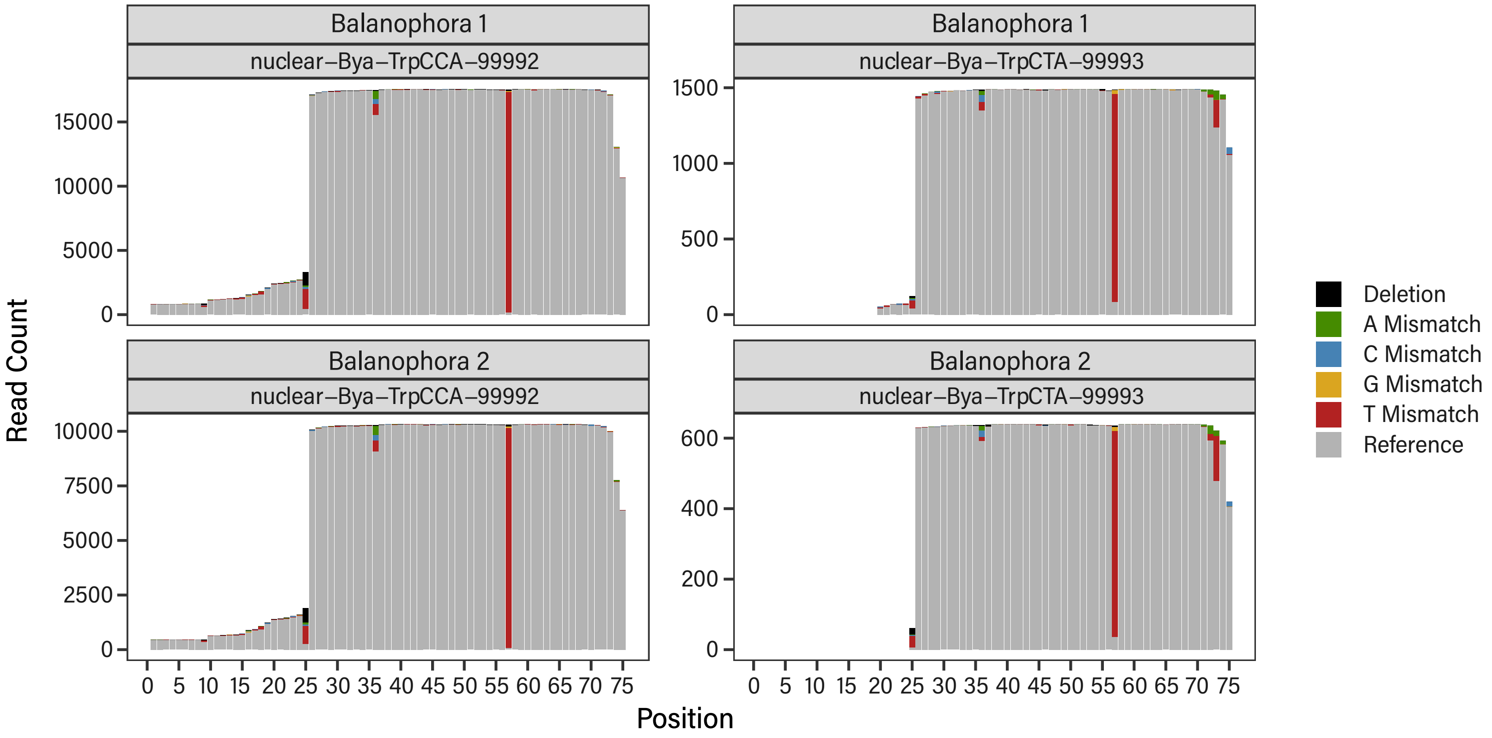

Figure S5. Read coverage and sequence variants in nuclear Trp tRNAs in *Lophophytum pyramidale*, and *Ombrophytum subterraneum*. Read counts were summed across replicates for each species after excluding reads that are truncated at the 3′ end (lacking >7 nt). Some variants, particularly those approaching 100% read coverage, may reflect differences between the genomic DNA sequences of Arabidopsis and the parasites rather than post-transcriptional modifications.

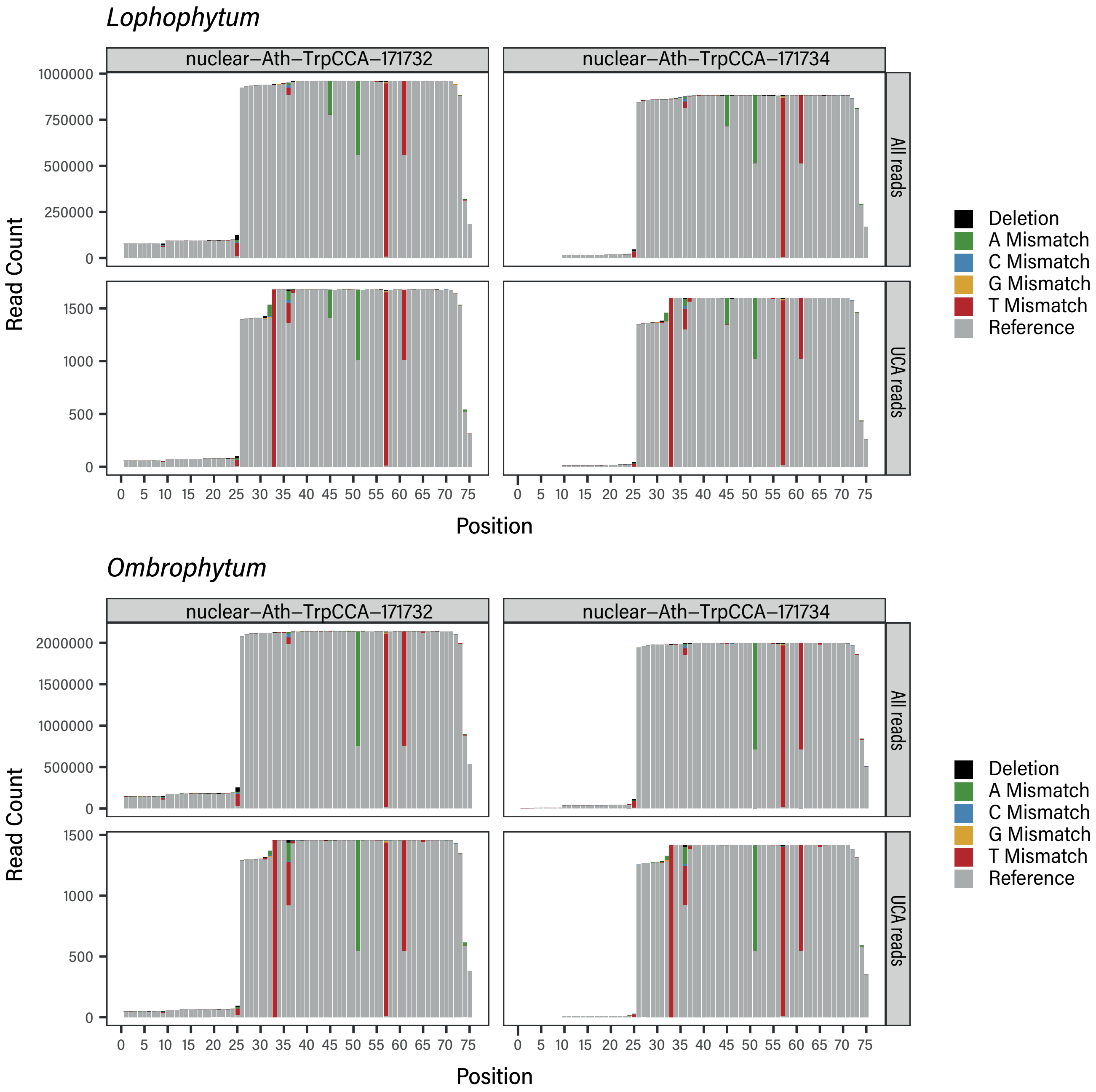

**Figure S6.** Maximum likelihood phylogenetic trees of aminoacyl-tRNA synthetases (aaRSs) from Balanophoraceae. **A.** Trees providing evidence of foreign aaRSs in Balanophoraceae (in red font). **B.** Trees showing the native origin of aaRSs that interact with functional foreign organellar-encoded tRNAs. Bootstrap support values >50% are shown above the branches. Branch lengths indicate substitutions per site.

**A**

cytosolic ThrRS

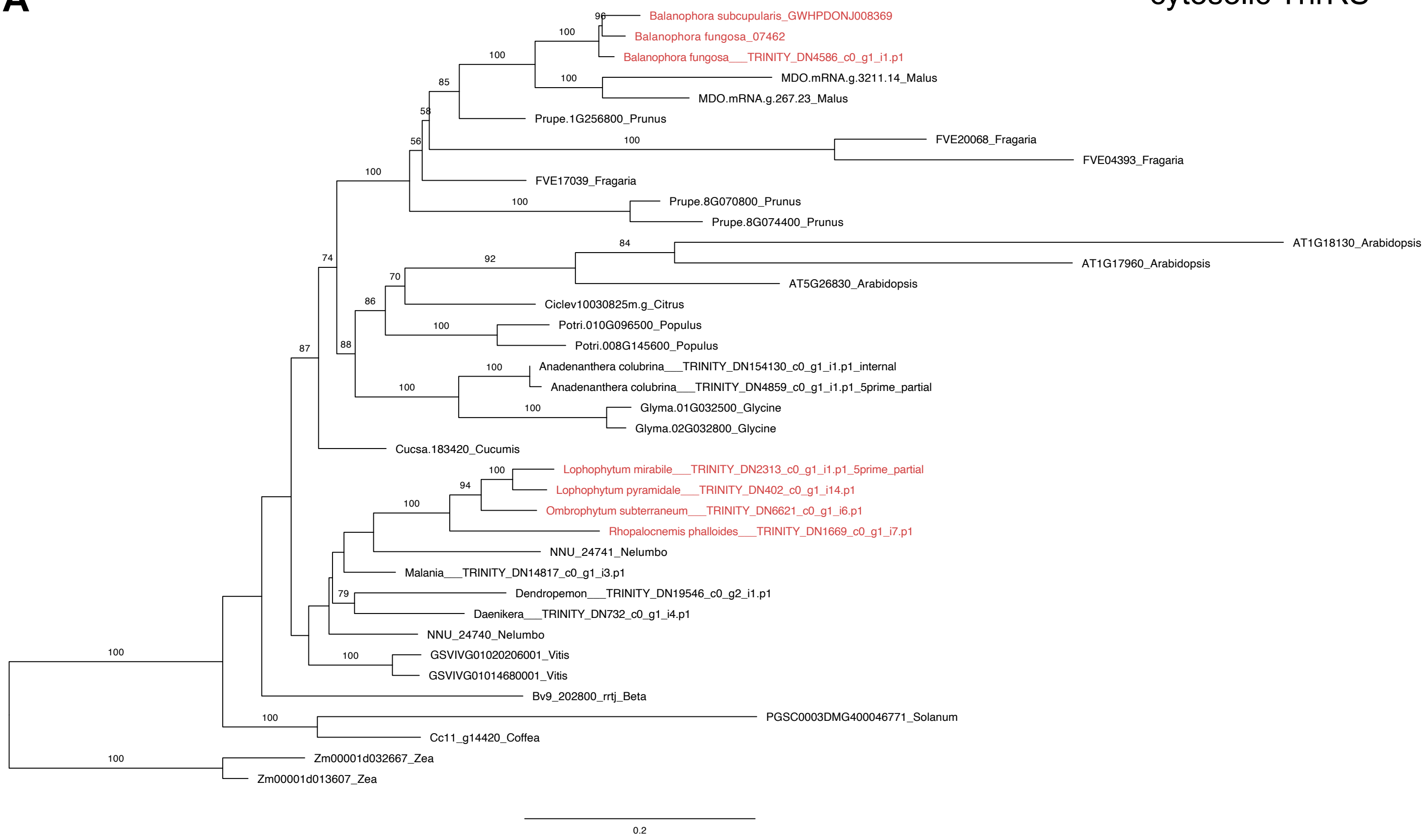


cytosolic TyrRS

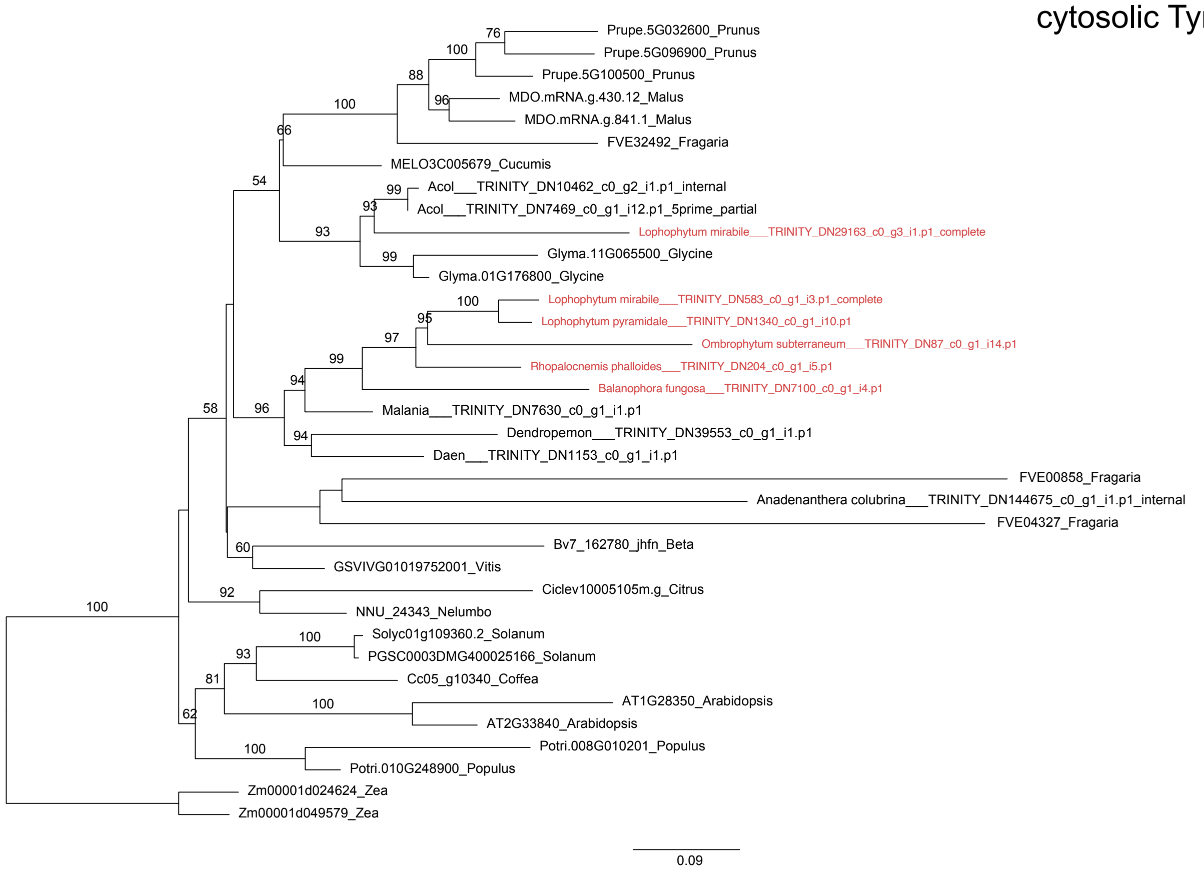

**B**

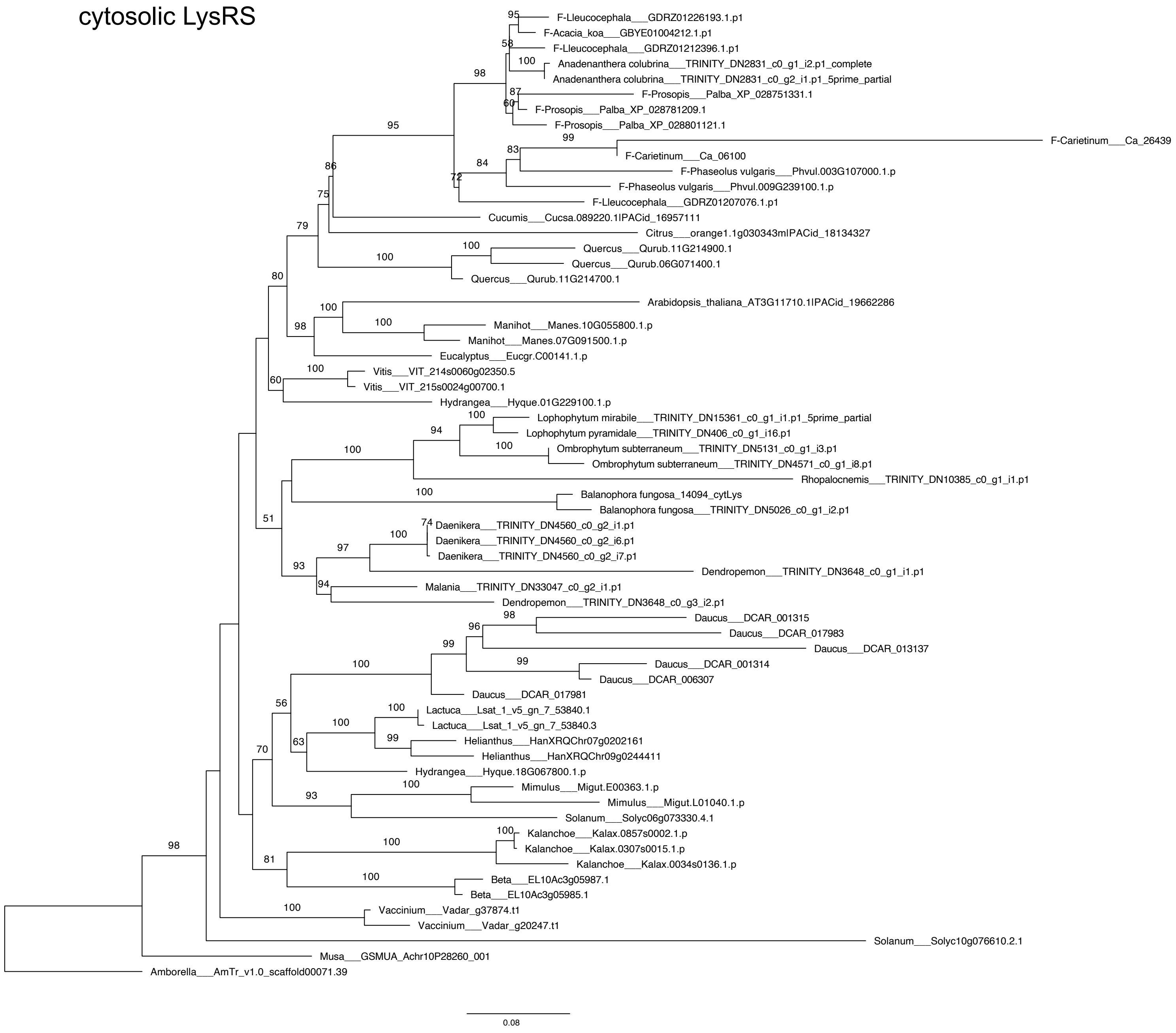
cytosolic LysRS
F-Lleucocephala___GDRZ01226193.1.p1
F-Acacia_koa___GBYE01004212.1.p1
F-Lleucocephala___GDRZ01212396.1.p1
Anadenanthera colubrina___TRINITY_DN2831_c0_g1_i2.p1_complete
Anadenanthera colubrina___TRINITY_DN2831_c0_g2_i1.p1_5prime_partial
F-Prosopis___Palba_XP_028751331.1
F-Prosopis___Palba_XP_028781209.1
F-Prosopis___Palba_XP_028801121.1
F-Carietinum___Ca_26439
F-Carietinum___Ca_06100
F-Phaseolus vulgaris___Phvul.003G107000.1.p
F-Phaseolus vulgaris___Phvul.009G239100.1.p
F-Lleucocephala___GDRZ01207076.1.p1
Cucumis___Cucsa.089220.1IPACid_16957111
Citrus___orange1.1g030343mIPACid_18134327
Quercus___Qurub.11G214900.1
Quercus___Qurub.06G071400.1
Quercus___Qurub.11G214700.1
Arabidopsis_thaliana_AT3G11710.1IPACid_19662286
Manihot___Manes.10G055800.1.p
Manihot___Manes.07G091500.1.p
Eucalyptus___Eucgr.C00141.1.p
Vitis___VIT_214s0060g02350.5
Vitis___VIT_215s0024g00700.1
Hydrangea___Hyque.01G229100.1.p
Lophophytum mirabile___TRINITY_DN15361_c0_g1_i1.p1_5prime_partial
Lophophytum pyramidale___TRINITY_DN406_c0_g1_i16.p1
Ombrophytum subterraneum___TRINITY_DN5131_c0_g1_i3.p1
Ombrophytum subterraneum___TRINITY_DN4571_c0_g1_i8.p1
Rhopalocnemis___TRINITY_DN10385_c0_g1_i1.p1
Balanophora fungosa_14094_cytLys
Balanophora fungosa___TRINITY_DN5026_c0_g1_i2.p1
Daenikera___TRINITY_DN4560_c0_g2_i1.p1
Daenikera___TRINITY_DN4560_c0_g2_i6.p1
Daenikera___TRINITY_DN4560_c0_g2_i7.p1
Dendropemon___TRINITY_DN3648_c0_g1_i1.p1
Malania___TRINITY_DN33047_c0_g2_i1.p1
Dendropemon___TRINITY_DN3648_c0_g3_i2.p1
Daucus___DCAR_001315
Daucus___DCAR_017983
Daucus___DCAR_013137
Daucus___DCAR_001314
Daucus___DCAR_006307
Daucus___DCAR_017981
Lactuca___Lsat_1_v5_gn_7_53840.1
Lactuca___Lsat_1_v5_gn_7_53840.3
Helianthus___HanXRQChr07g0202161
Helianthus___HanXRQChr09g0244411
Hydrangea___Hyque.18G067800.1.p
Mimulus___Migut.E00363.1.p
Mimulus___Migut.L01040.1.p
Solanum___Solyc06g073330.4.1
Kalanchoe___Kalax.0857s0002.1.p
Kalanchoe___Kalax.0307s0015.1.p
Kalanchoe___Kalax.0034s0136.1.p
Beta___EL10Ac3g05987.1
Beta___EL10Ac3g05985.1
Vaccinium___Vadar_g37874.t1
Vaccinium___Vadar_g20247.t1
Solanum___Solyc10g076610.2.1
Musa___GSMUA_Achr10P28260_001
Amborella___AmTr_v1.0_scaffold00071.39
0.08

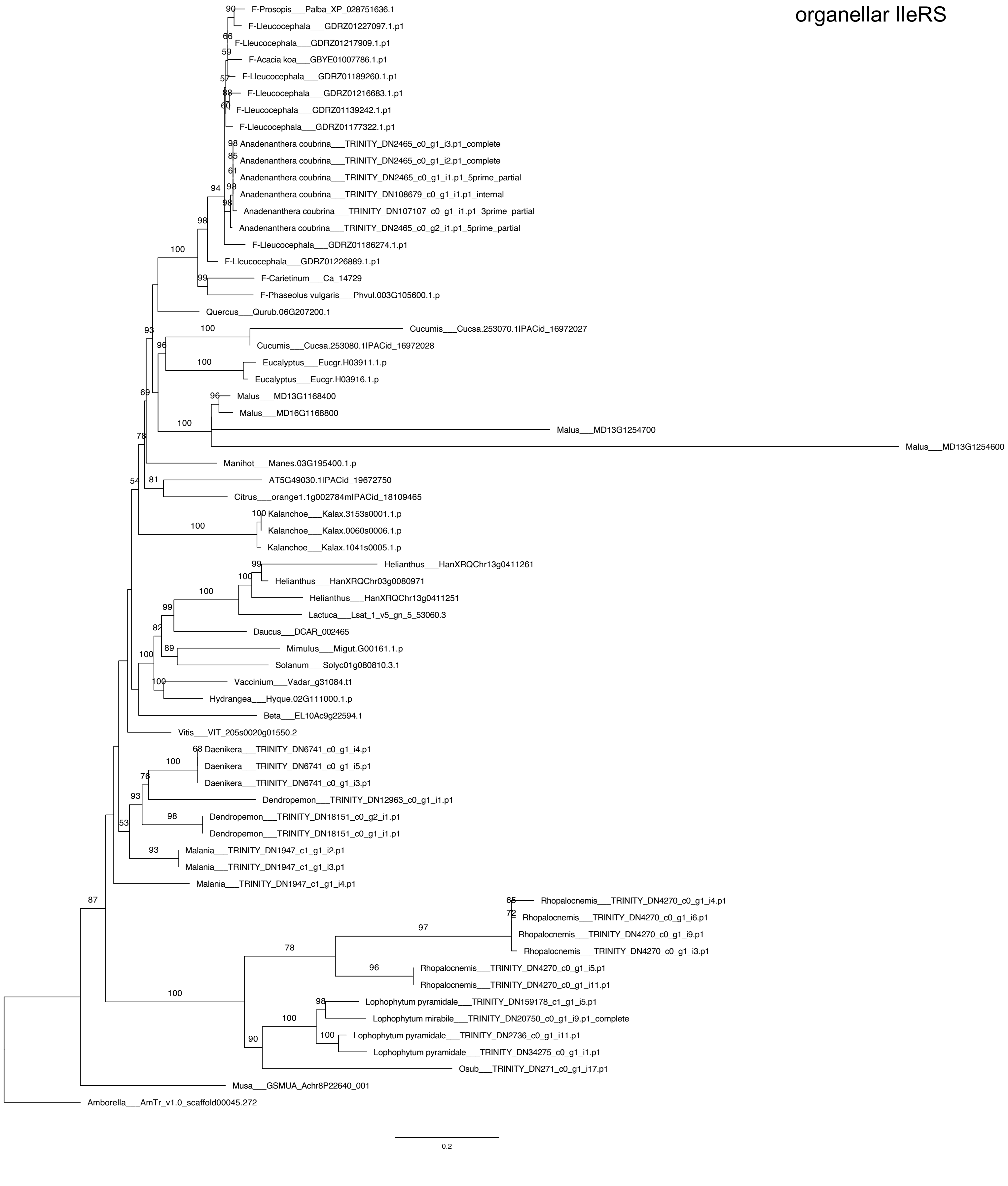

organellar IleRS
F-Prosopis___Palba_XP_028751636.1
F-Lleucocephala___GDRZ01227097.1.p1
F-Lleucocephala___GDRZ01217909.1.p1
F-Acacia koa___GBYE01007786.1.p1
F-Lleucocephala___GDRZ01189260.1.p1
F-Lleucocephala___GDRZ01216683.1.p1
F-Lleucocephala___GDRZ01139242.1.p1
F-Lleucocephala___GDRZ01177322.1.p1
Anadenanthera coubrina___TRINITY_DN2465_c0_g1_i3.p1_complete
Anadenanthera coubrina___TRINITY_DN2465_c0_g1_i2.p1_complete
Anadenanthera coubrina___TRINITY_DN2465_c0_g1_i1.p1_5prime_partial
Anadenanthera coubrina___TRINITY_DN108679_c0_g1_i1.p1_internal
Anadenanthera coubrina___TRINITY_DN107107_c0_g1_i1.p1_3prime_partial
Anadenanthera coubrina___TRINITY_DN2465_c0_g2_i1.p1_5prime_partial
F-Lleucocephala___GDRZ01186274.1.p1
F-Lleucocephala___GDRZ01226889.1.p1
F-Carietinum___Ca_14729
F-Phaseolus vulgaris___Phvul.003G105600.1.p
Quercus___Qurub.06G207200.1
Cucumis___Cucsa.253070.1IPACid_16972027
Cucumis___Cucsa.253080.1IPACid_16972028
Eucalyptus___Eucgr.H03911.1.p
Eucalyptus___Eucgr.H03916.1.p
Malus___MD13G1168400
Malus___MD16G1168800
Malus___MD13G1254700
Malus___MD13G1254600
Manihot___Manes.03G195400.1.p
AT5G49030.1IPACid_19672750
Citrus___orange1.1g002784mIPACid_18109465
Kalanchoe___Kalax.3153s0001.1.p
Kalanchoe___Kalax.0060s0006.1.p
Kalanchoe___Kalax.1041s0005.1.p
Helianthus___HanXRQChr13g0411261
Helianthus___HanXRQChr03g0080971
Helianthus___HanXRQChr13g0411251
Lactuca___Lsat_1_v5_gn_5_53060.3
Daucus___DCAR_002465
Mimulus___Migut.G00161.1.p
Solanum___Solyc01g080810.3.1
Vaccinium___Vadar_g31084.t1
Hydrangea___Hyque.02G111000.1.p
Beta___EL10Ac9g22594.1
Vitis___VIT_205s0020g01550.2
Daenikera___TRINITY_DN6741_c0_g1_i4.p1
Daenikera___TRINITY_DN6741_c0_g1_i5.p1
Daenikera___TRINITY_DN6741_c0_g1_i3.p1
Dendropemon___TRINITY_DN12963_c0_g1_i1.p1
Dendropemon___TRINITY_DN18151_c0_g2_i1.p1
Dendropemon___TRINITY_DN18151_c0_g1_i1.p1
Malania___TRINITY_DN1947_c1_g1_i2.p1
Malania___TRINITY_DN1947_c1_g1_i3.p1
Malania___TRINITY_DN1947_c1_g1_i4.p1
Rhopalocnemis___TRINITY_DN4270_c0_g1_i4.p1
Rhopalocnemis___TRINITY_DN4270_c0_g1_i6.p1
Rhopalocnemis___TRINITY_DN4270_c0_g1_i9.p1
Rhopalocnemis___TRINITY_DN4270_c0_g1_i3.p1
Rhopalocnemis___TRINITY_DN4270_c0_g1_i5.p1
Rhopalocnemis___TRINITY_DN4270_c0_g1_i11.p1
Lophophytum pyramidale___TRINITY_DN159178_c1_g1_i5.p1
Lophophytum mirabile___TRINITY_DN20750_c0_g1_i9.p1_complete
Lophophytum pyramidale___TRINITY_DN2736_c0_g1_i11.p1
Lophophytum pyramidale___TRINITY_DN34275_c0_g1_i1.p1
Osub___TRINITY_DN271_c0_g1_i17.p1
Musa___GSMUA_Achr8P22640_001
Amborella___AmTr_v1.0_scaffold00045.272
0.2

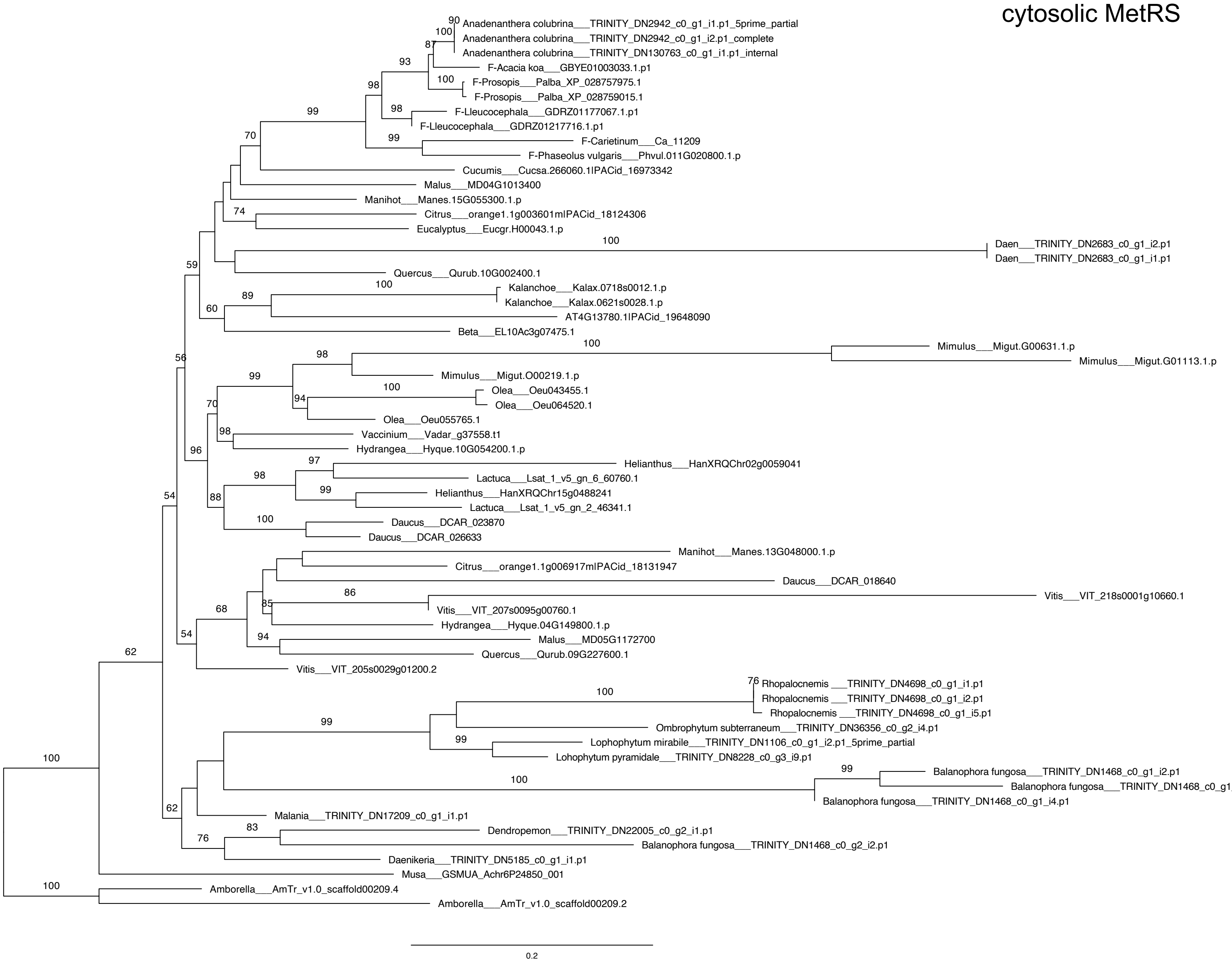
cytosolic MetRS
Anadenanthera colubrina___TRINITY_DN2942_c0_g1_i1.p1_5prime_partial
Anadenanthera colubrina___TRINITY_DN2942_c0_g1_i2.p1_complete
Anadenanthera colubrina___TRINITY_DN130763_c0_g1_i1.p1_internal
F-Acacia koa___GBYE01003033.1.p1
F-Prosopis___Palba_XP_028757975.1
F-Prosopis___Palba_XP_028759015.1
F-Lleucocephala___GDRZ01177067.1.p1
F-Lleucocephala___GDRZ01217716.1.p1
F-Carietinum___Ca_11209
F-Phaseolus vulgaris___Phvul.011G020800.1.p
Cucumis___Cucsa.266060.1lPACid_16973342
Malus___MD04G1013400
Manihot___Manes.15G055300.1.p
Citrus___orange1.1g003601mlPACid_18124306
Eucalyptus___Eucgr.H00043.1.p
Daen___TRINITY_DN2683_c0_g1_i2.p1
Daen___TRINITY_DN2683_c0_g1_i1.p1
Quercus___Qurub.10G002400.1
Kalanchoe___Kalax.0718s0012.1.p
Kalanchoe___Kalax.0621s0028.1.p
AT4G13780.1lPACid_19648090
Beta___EL10Ac3g07475.1
Mimulus___Migut.G00631.1.p
Mimulus___Migut.G01113.1.p
Mimulus___Migut.O00219.1.p
Olea___Oeu043455.1
Olea___Oeu064520.1
Olea___Oeu055765.1
Vaccinium___Vadar_g37558.t1
Hydrangea___Hyque.10G054200.1.p
Helianthus___HanXRQChr02g0059041
Lactuca___Lsat_1_v5_gn_6_60760.1
Helianthus___HanXRQChr15g0488241
Lactuca___Lsat_1_v5_gn_2_46341.1
Daucus___DCAR_023870
Daucus___DCAR_026633
Manihot___Manes.13G048000.1.p
Citrus___orange1.1g006917mlPACid_18131947
Daucus___DCAR_018640
Vitis___VIT_218s0001g10660.1
Vitis___VIT_207s0095g00760.1
Hydrangea___Hyque.04G149800.1.p
Malus___MD05G1172700
Quercus___Qurub.09G227600.1
Vitis___VIT_205s0029g01200.2
Rhopalocnemis ___TRINITY_DN4698_c0_g1_i1.p1
Rhopalocnemis ___TRINITY_DN4698_c0_g1_i2.p1
Rhopalocnemis ___TRINITY_DN4698_c0_g1_i5.p1
Ombrophytum subterraneum___TRINITY_DN36356_c0_g2_i4.p1
Lophophytum mirabile___TRINITY_DN1106_c0_g1_i2.p1_5prime_partial
Lohophytum pyramidale___TRINITY_DN8228_c0_g3_i9.p1
Balanophora fungosa___TRINITY_DN1468_c0_g1_i2.p1
Balanophora fungosa___TRINITY_DN1468_c0_g
Balanophora fungosa___TRINITY_DN1468_c0_g1_i4.p1
Malania___TRINITY_DN17209_c0_g1_i1.p1
Dendropemon___TRINITY_DN22005_c0_g2_i1.p1
Balanophora fungosa___TRINITY_DN1468_c0_g2_i2.p1
Daenikeria___TRINITY_DN5185_c0_g1_i1.p1
Musa___GSMUA_Achr6P24850_001
Amborella___AmTr_v1.0_scaffold00209.4
Amborella___AmTr_v1.0_scaffold00209.2
0.2

### cytosolic GluRS

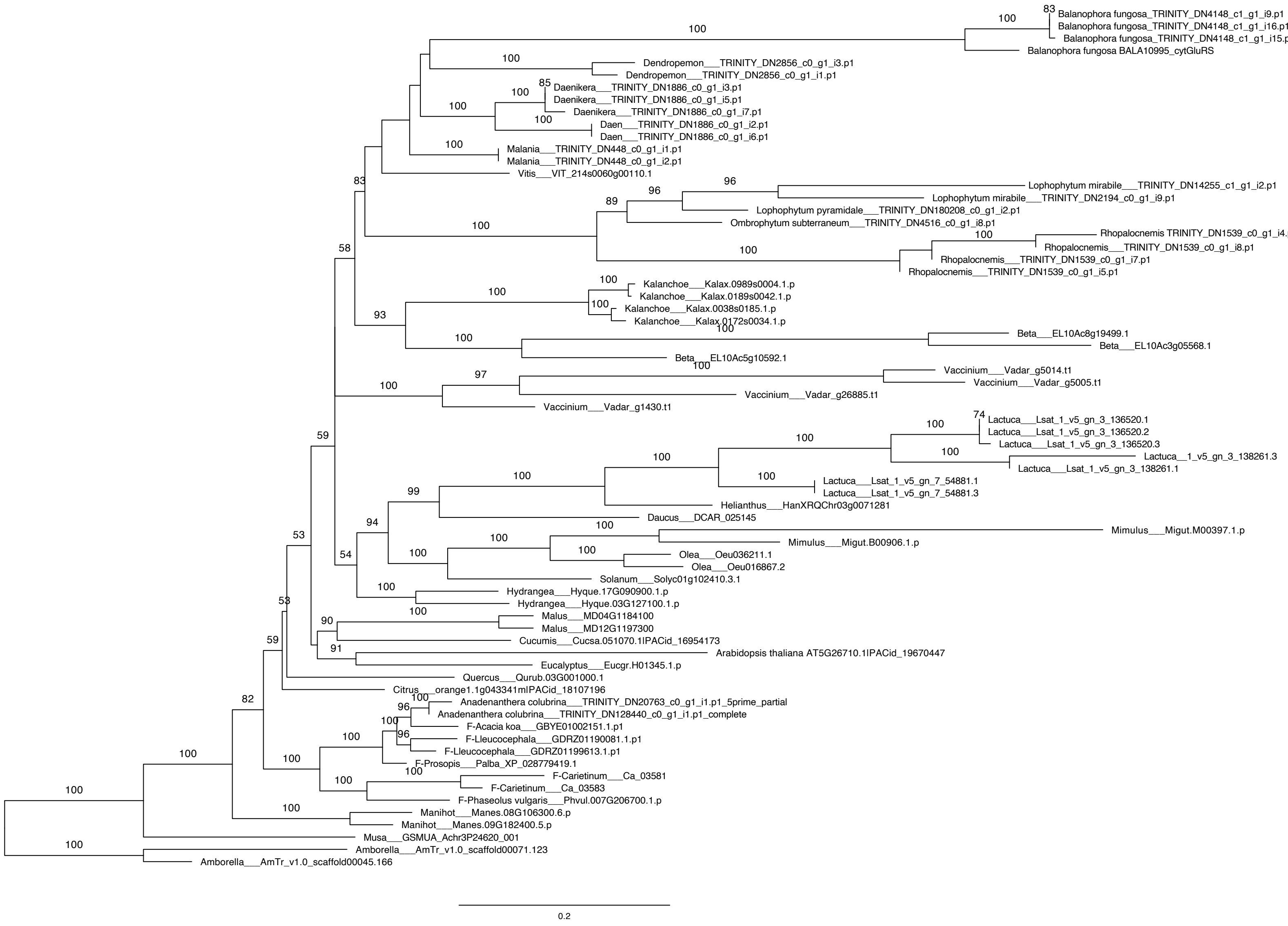

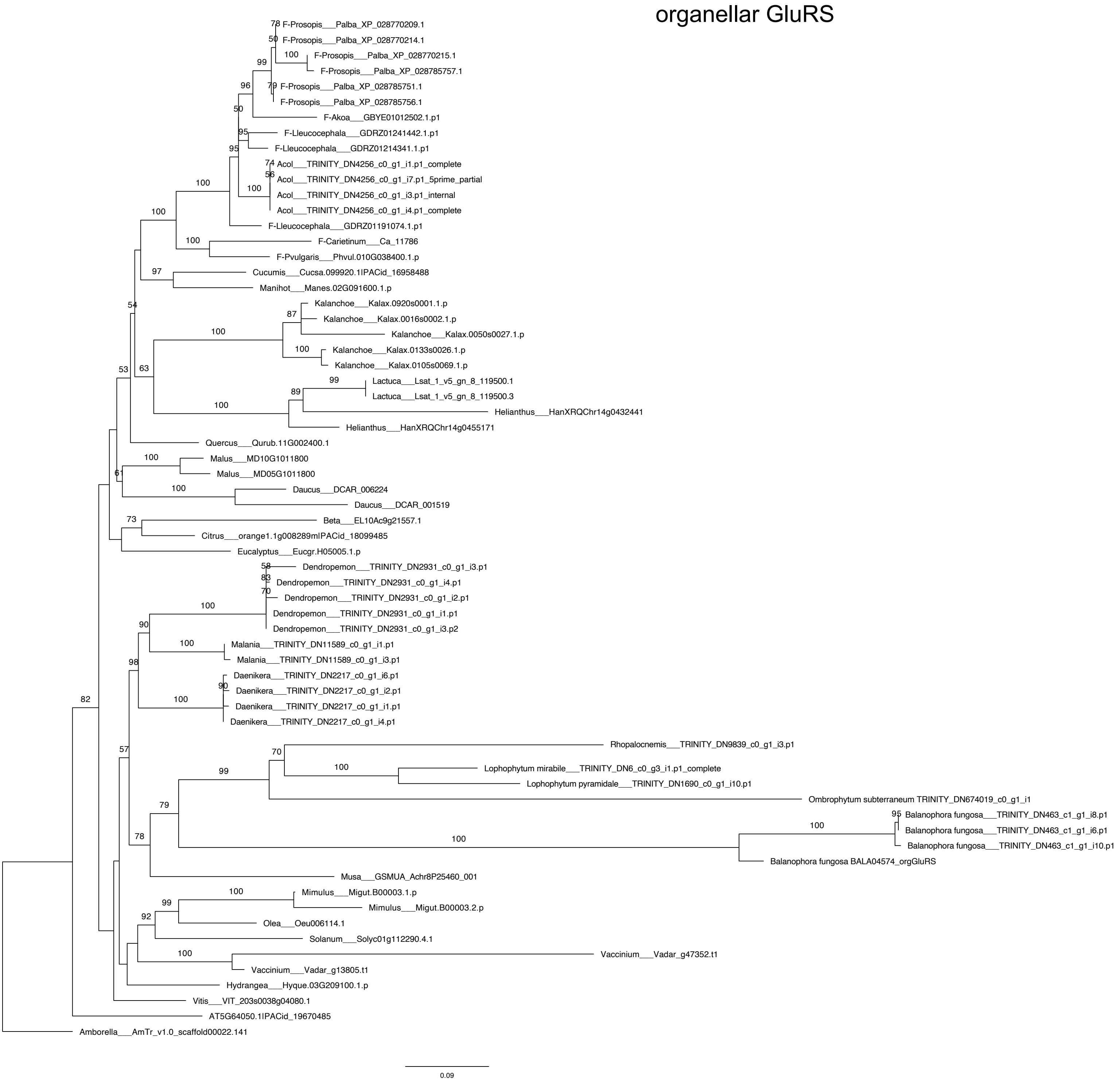
organellar GluRS
F-Prosopis___Palba_XP_028770209.1
F-Prosopis___Palba_XP_028770214.1
F-Prosopis___Palba_XP_028770215.1
F-Prosopis___Palba_XP_028785757.1
F-Prosopis___Palba_XP_028785751.1
F-Prosopis___Palba_XP_028785756.1
F-Akoa___GBYE01012502.1.p1
F-Lleucocephala___GDRZ01241442.1.p1
F-Lleucocephala___GDRZ01214341.1.p1
Acol___TRINITY_DN4256_c0_g1_i1.p1_complete
Acol___TRINITY_DN4256_c0_g1_i7.p1_5prime_partial
Acol___TRINITY_DN4256_c0_g1_i3.p1_internal
Acol___TRINITY_DN4256_c0_g1_i4.p1_complete
F-Lleucocephala___GDRZ01191074.1.p1
F-Carietinum___Ca_11786
F-Pvulgaris___Phvul.010G038400.1.p
Cucumis___Cucsa.099920.1IPACid_16958488
Manihot___Manes.02G091600.1.p
Kalanchoe___Kalax.0920s0001.1.p
Kalanchoe___Kalax.0016s0002.1.p
Kalanchoe___Kalax.0050s0027.1.p
Kalanchoe___Kalax.0133s0026.1.p
Kalanchoe___Kalax.0105s0069.1.p
Lactuca___Lsat_1_v5_gn_8_119500.1
Lactuca___Lsat_1_v5_gn_8_119500.3
Helianthus___HanXRQChr14g0432441
Helianthus___HanXRQChr14g0455171
Quercus___Qurub.11G002400.1
Malus___MD10G1011800
Malus___MD05G1011800
Daucus___DCAR_006224
Daucus___DCAR_001519
Beta___EL10Ac9g21557.1
Citrus___orange1.1g008289mIPACid_18099485
Eucalyptus___Eucgr.H05005.1.p
Dendropemon___TRINITY_DN2931_c0_g1_i3.p1
Dendropemon___TRINITY_DN2931_c0_g1_i4.p1
Dendropemon___TRINITY_DN2931_c0_g1_i2.p1
Dendropemon___TRINITY_DN2931_c0_g1_i1.p1
Dendropemon___TRINITY_DN2931_c0_g1_i3.p2
Malania___TRINITY_DN11589_c0_g1_i1.p1
Malania___TRINITY_DN11589_c0_g1_i3.p1
Daenikera___TRINITY_DN2217_c0_g1_i6.p1
Daenikera___TRINITY_DN2217_c0_g1_i2.p1
Daenikera___TRINITY_DN2217_c0_g1_i1.p1
Daenikera___TRINITY_DN2217_c0_g1_i4.p1
Rhopalocnemis___TRINITY_DN9839_c0_g1_i3.p1
Lophophytum mirabile___TRINITY_DN6_c0_g3_i1.p1_complete
Lophophytum pyramidale___TRINITY_DN1690_c0_g1_i10.p1
Ombrophytum subterraneum TRINITY_DN674019_c0_g1_i1
Balanophora fungosa___TRINITY_DN463_c1_g1_i8.p1
Balanophora fungosa___TRINITY_DN463_c1_g1_i6.p1
Balanophora fungosa___TRINITY_DN463_c1_g1_i10.p1
Balanophora fungosa BALA04574_orgGluRS
Musa___GSMUA_Achr8P25460_001
Mimulus___Migut.B00003.1.p
Mimulus___Migut.B00003.2.p
Olea___Oeu006114.1
Solanum___Solyc01g112290.4.1
Vaccinium___Vadar_g47352.t1
Vaccinium___Vadar_g13805.t1
Hydrangea___Hyque.03G209100.1.p
Vitis___VIT_203s0038g04080.1
AT5G64050.1IPACid_19670485
Amborella___AmTr_v1.0_scaffold00022.141
0.09

MTF

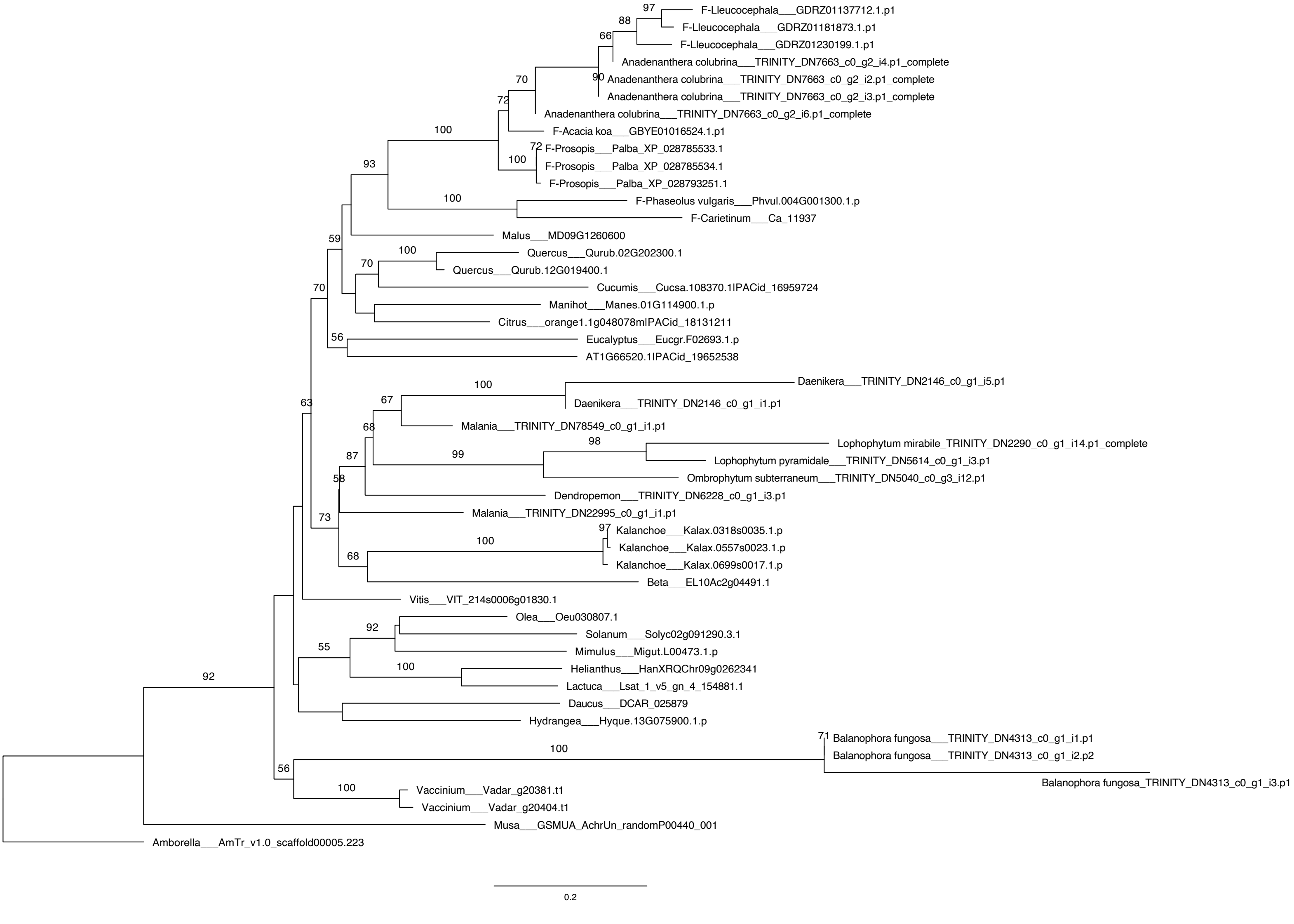
97
88
66
F-Lleucocephala___GDRZ01137712.1.p1
F-Lleucocephala___GDRZ01181873.1.p1
F-Lleucocephala___GDRZ01230199.1.p1
Anadenanthera colubrina___TRINITY_DN7663_c0_g2_i4.p1_complete
90
Anadenanthera colubrina___TRINITY_DN7663_c0_g2_i2.p1_complete
Anadenanthera colubrina___TRINITY_DN7663_c0_g2_i3.p1_complete
70
72
Anadenanthera colubrina___TRINITY_DN7663_c0_g2_i6.p1_complete
100
F-Acacia koa___GBYE01016524.1.p1
72
F-Prosopis___Palba_XP_028785533.1
100
F-Prosopis___Palba_XP_028785534.1
F-Prosopis___Palba_XP_028793251.1
93
100
F-Phaseolus vulgaris___Phvul.004G001300.1.p
F-Carietinum___Ca_11937
Malus___MD09G1260600
59
100
Quercus___Qurub.02G202300.1
70
Quercus___Qurub.12G019400.1
70
Cucumis___Cucsa.108370.1lPACid_16959724
Manihot___Manes.01G114900.1.p
Citrus___orange1.1g048078mlPACid_18131211
56
Eucalyptus___Eucgr.F02693.1.p
AT1G66520.1lPACid_19652538
100
Daenikera___TRINITY_DN2146_c0_g1_i5.p1
63
67
Daenikera___TRINITY_DN2146_c0_g1_i1.p1
68
Malania___TRINITY_DN78549_c0_g1_i1.p1
98
Lophophytum mirabile_TRINITY_DN2290_c0_g1_i14.p1_complete
87
99
Lophophytum pyramidale___TRINITY_DN5614_c0_g1_i3.p1
58
Ombrophytum subterraneum___TRINITY_DN5040_c0_g3_i12.p1
Dendropemon___TRINITY_DN6228_c0_g1_i3.p1
73
Malania___TRINITY_DN22995_c0_g1_i1.p1
97
Kalanchoe___Kalax.0318s0035.1.p
100
Kalanchoe___Kalax.0557s0023.1.p
68
Kalanchoe___Kalax.0699s0017.1.p
Beta___EL10Ac2g04491.1
Vitis___VIT_214s0006g01830.1
Olea___Oeu030807.1
92
Solanum___Solyc02g091290.3.1
55
Mimulus___Migut.L00473.1.p
100
Helianthus___HanXRQChr09g0262341
92
Lactuca___Lsat_1_v5_gn_4_154881.1
Daucus___DCAR_025879
Hydrangea___Hyque.13G075900.1.p
71
Balanophora fungosa___TRINITY_DN4313_c0_g1_i1.p1
100
Balanophora fungosa___TRINITY_DN4313_c0_g1_i2.p2
56
Balanophora fungosa_TRINITY_DN4313_c0_g1_i3.p1
100
Vaccinium___Vadar_g20381.t1
Vaccinium___Vadar_g20404.t1
Musa___GSMUA_AchrUn_randomP00440_001
Amborella___AmTr_v1.0_scaffold00005.223
0.2

### cytosolic TrpRS

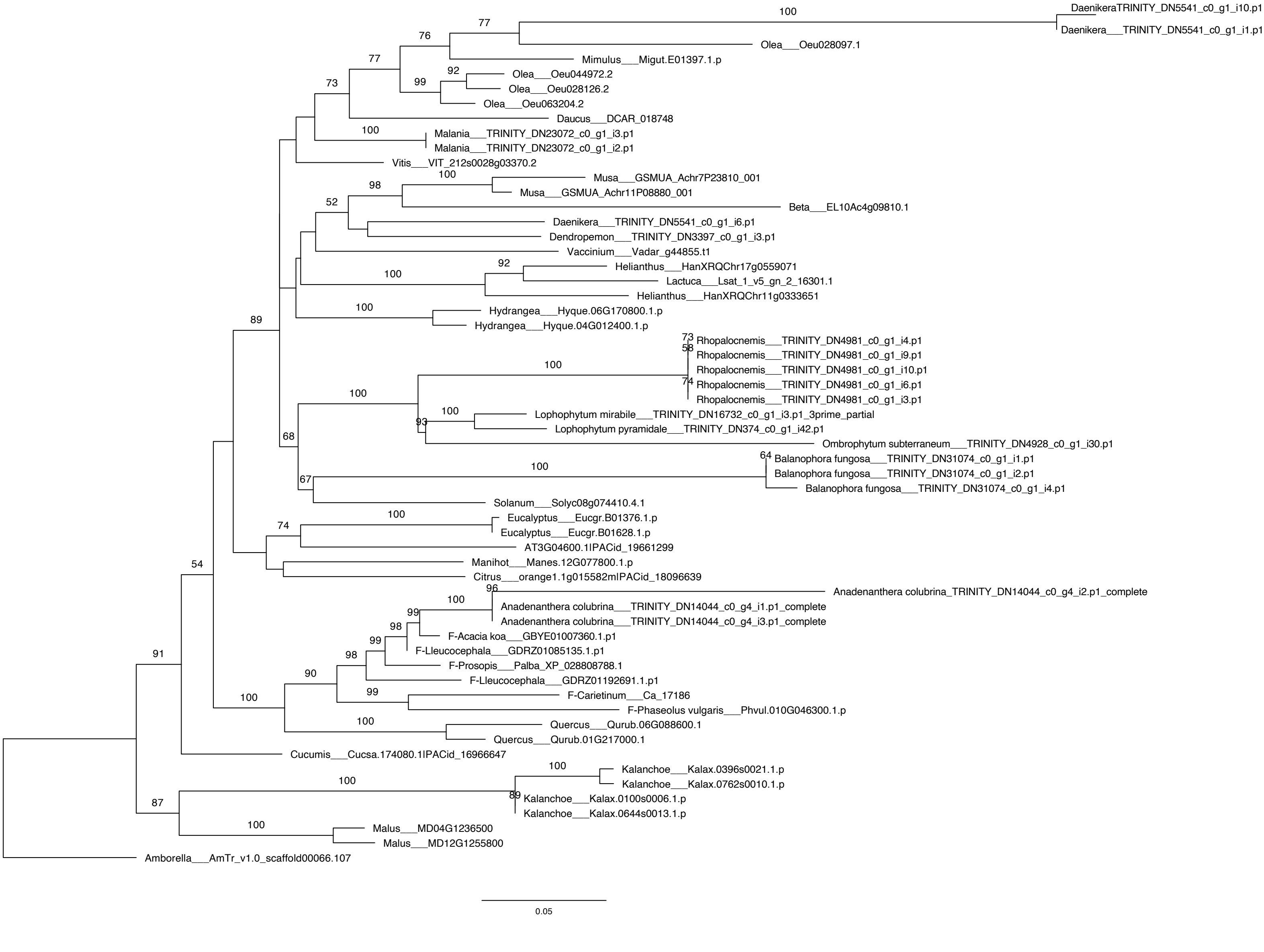

### organellar TrpRS

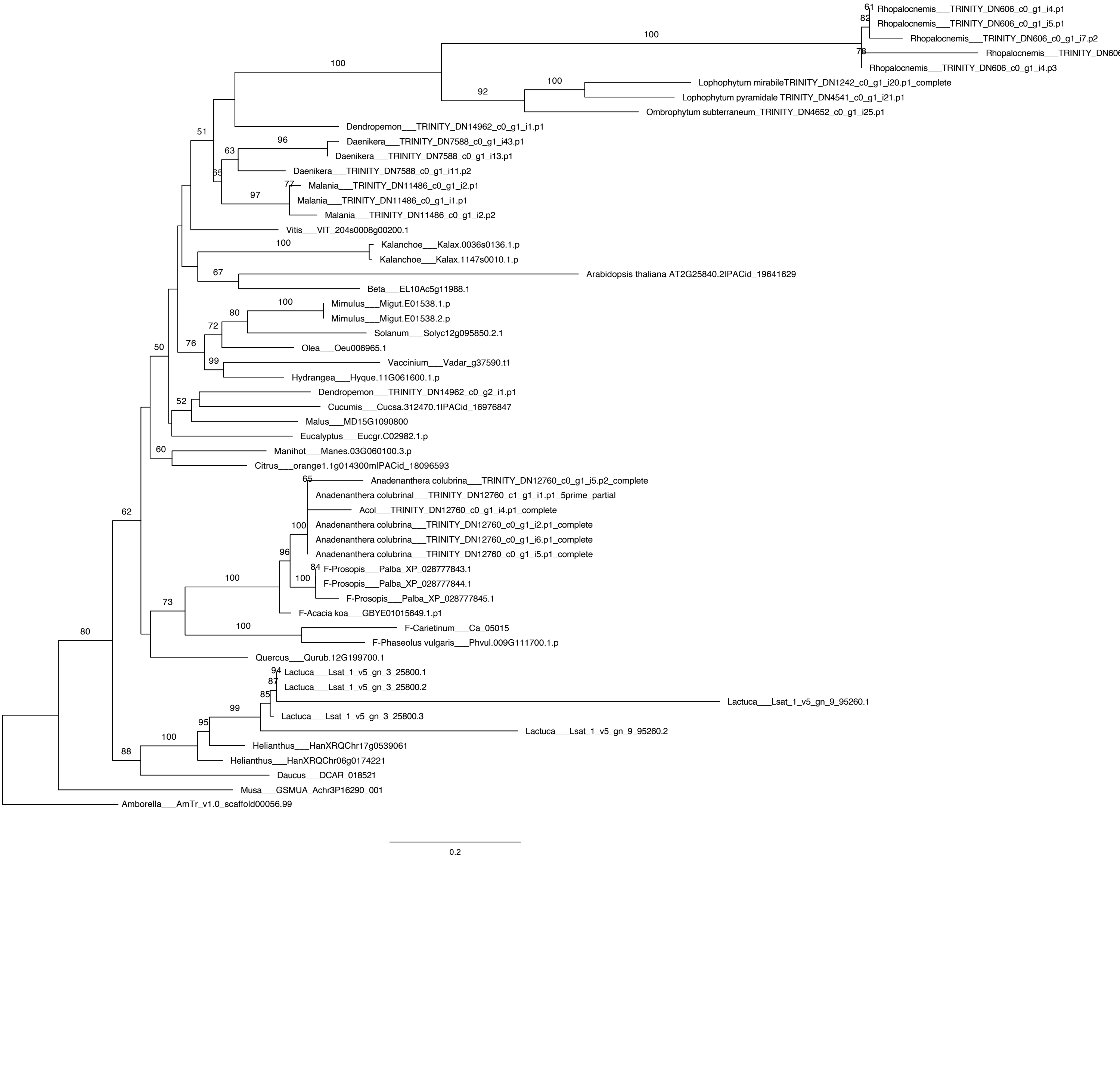

**Figure S7**. Transit peptides from Balanophoraceae target organellar GluRS to plastids. **A.** Fluorescent confocal images of localization for GFP fused to N-terminal transit peptides from organellar-type GluRS from three different species. Chlorophyll autofluorescence and IVD-FP611 are used to visualize plastids and mitochondria, respectively. **B.** No-GFP control (i.e. just IVD-FP611) visualized under the same settings and conditions as test constructs. **C.** Additional images of localization for GFP construct with an N-terminal transit peptide from cytosolic-type GluRS from *Lophophytum pyramidale*. Yellow brackets denote plastids exhibiting weak GFP localization. Note the lack of GFP signal for untransformed guard cell and pavement cell (outlined in yellow) compared to plastid from neighboring transformed cell (yellow bracket) showing true GFP fluorescence.

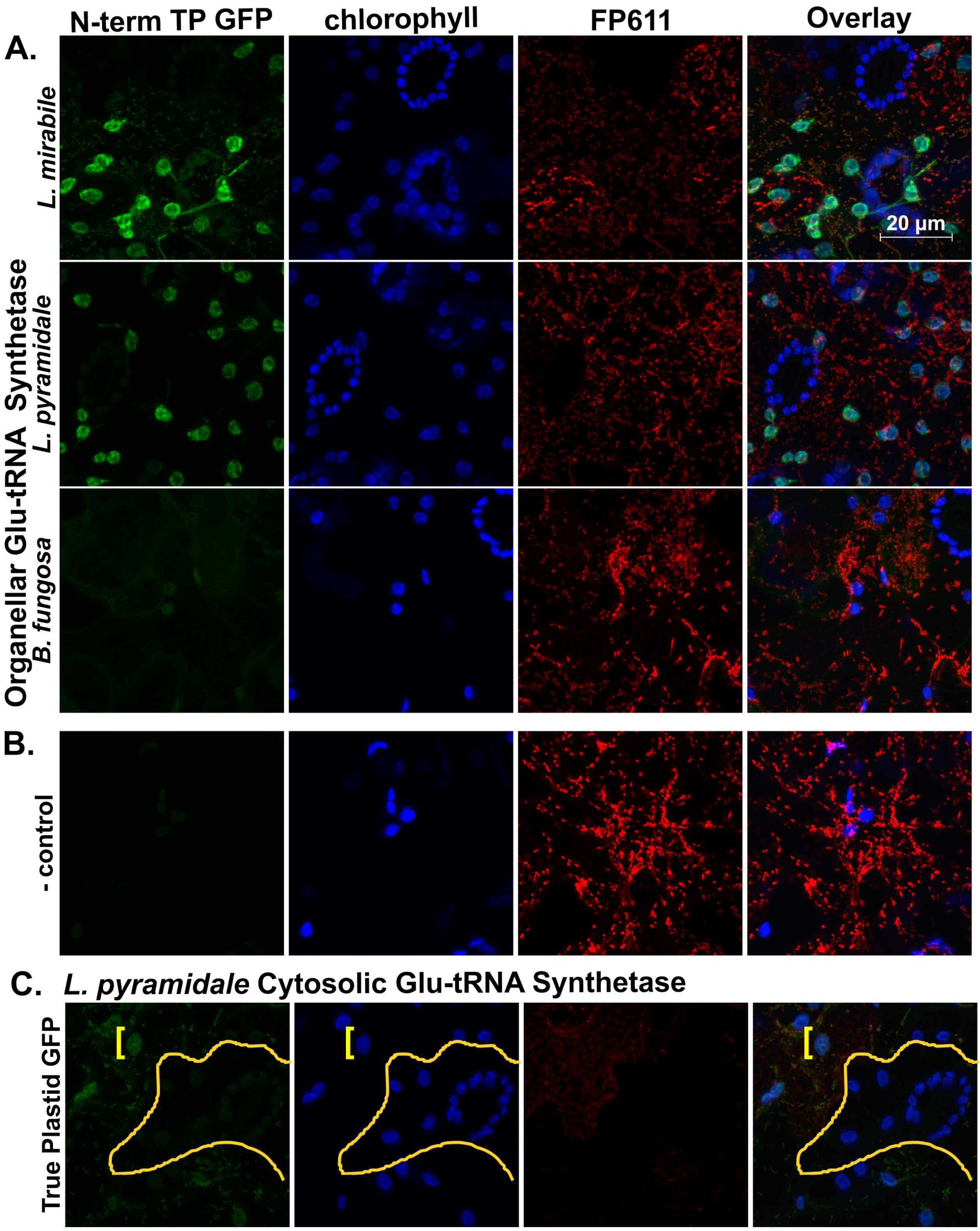

**Figure S8.** Subunit composition of plastid and mitochondrial ribosomes in Balanophoraceae and *Arabidopsis*, and identification of mitochondrial, plastid or nuclear-encoded genes.

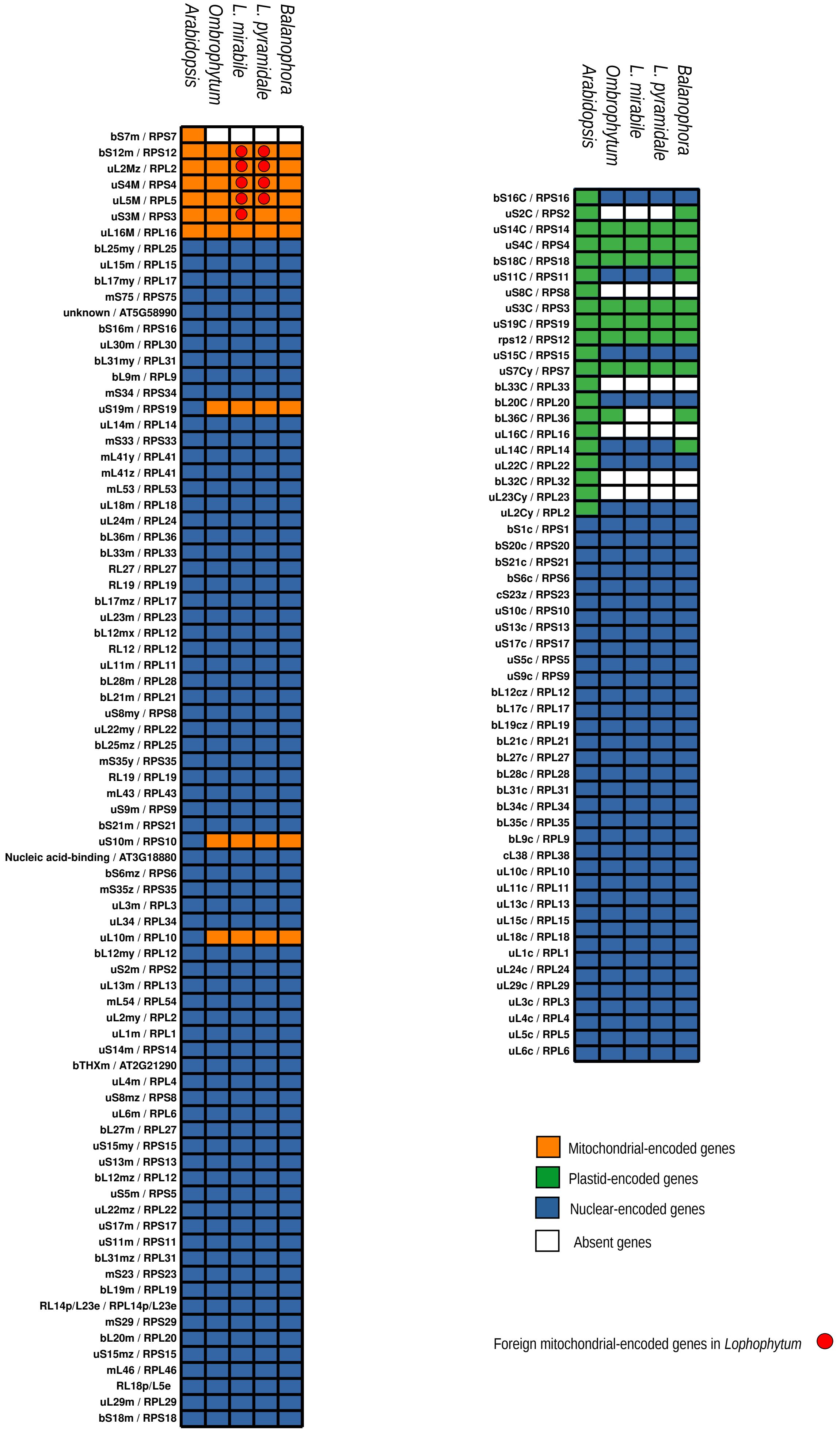

**Figure S9.** Evolutionary dynamics of organellar ribosomes in Balanophoraceae. Maximum likelihood phylogenies of two representative nuclear genes (*rps1* and *rpl23*) encoding plastid-targeted and mitochondrion-targeted ribosomal proteins, respectively. Sequences from Balanophoraceae are highlighted in blue. Conserved protein domains identified in *Balanophora fungosa* are indicated.

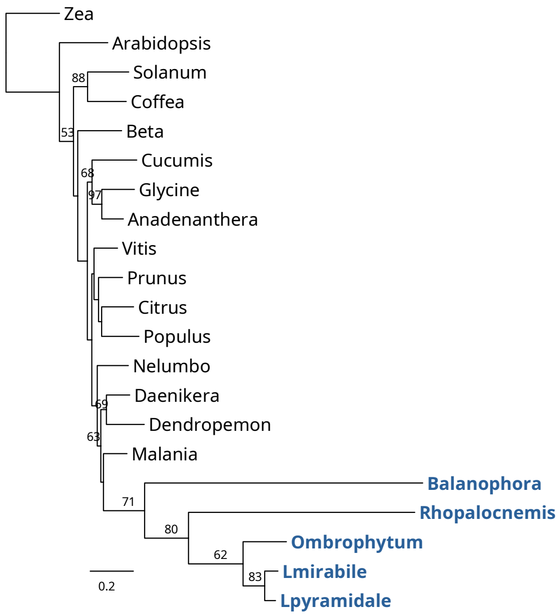


*rps1-Balanophora*

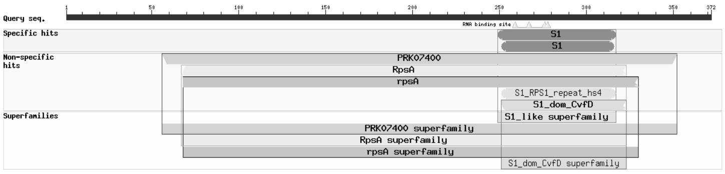


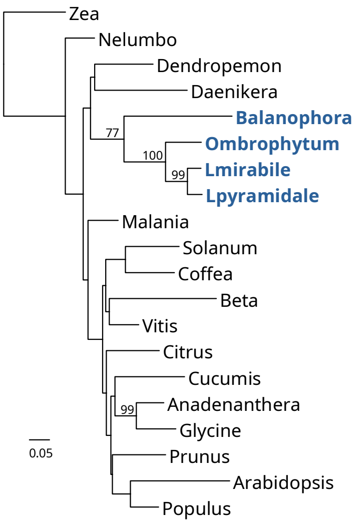


*rpl4-Balanophora*

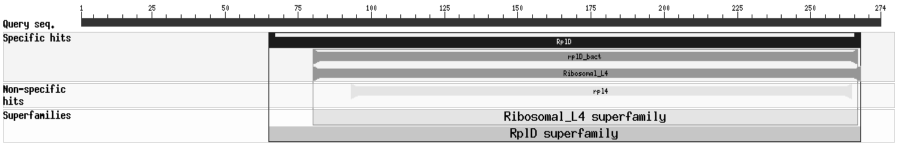
